# The trypanosome UDP-glucose pyrophosphorylase is imported by piggybacking into glycosomes where unconventional sugar nucleotide synthesis takes place

**DOI:** 10.1101/2021.02.17.431754

**Authors:** Oriana Villafraz, Hélène Baudouin, Muriel Mazet, Hanna Kulyk, Jean-William Dupuy, Erika Pineda, Cyrille Botté, Daniel Inaoka, Jean-Charles Portais, Frédéric Bringaud

## Abstract

Glycosomes are peroxisome-related organelles of trypanosomatid parasites containing metabolic pathways usually present in the cytosol of other eukaryotes, such as glycolysis and biosynthesis of sugar nucleotides. UDP-glucose pyrophosphorylase (UGP), the enzyme responsible for the synthesis of the sugar nucleotide UDP-glucose, is localised in the cytosol and glycosomes of the bloodstream and procyclic trypanosomes, despite the absence of any known peroxisomal targeting signal (PTS1 and PTS2). The questions we addressed here are (*i*) is the unusual glycosomal biosynthetic pathway of sugar nucleotide functional and (*ii*) how the PTS-free UGP is imported into glycosomes? We showed that UGP is imported into glycosomes by piggybacking on the glycosomal PTS1-containing phosphoenolpyruvate carboxykinase (PEPCK) and identified the domains involved in the UGP/PEPCK interaction. Proximity ligation assays revealed that this interaction occurs in 3-10% of glycosomes, suggesting that these correspond to organelles competent for protein import. We also showed that UGP is essential for growth of trypanosomes and that both the glycosomal and cytosolic metabolic pathways involving UGP are functional, since the lethality of the knock-down UGP mutant cell line (*^RNAi^*UGP) was rescued by expressing a recoded UGP in the organelle (*^RNAi^*UGP/*^EXP^*rUGP-GPDH). Our conclusion was supported by targeted metabolomic analyses (IC-HRMS) showing that UDP-glucose is no longer detectable in the *^RNAi^*UGP mutant, while it is still produced in cells expressing UGP exclusively in the cytosol (PEPCK null mutant) or glycosomes (*^RNAi^*UGP/*^EXP^*rUGP-GPDH). Trypanosomatids are the only known organisms to have selected functional peroxisomal (glycosomal) sugar nucleotide biosynthetic pathways in addition to the canonical cytosolic ones.

**Importance:** Unusual compartmentalization of metabolic pathways within organelles is one of the most enigmatic features of trypanosomatids. These unicellular eukaryotes are the only organisms that sequestered glycolysis inside peroxisomes (glycosomes), although the selective advantage of this compartmentalization is still not clear. Trypanosomatids are also unique for the glycosomal localisation of enzymes of the sugar nucleotide biosynthetic pathways, which are also present in the cytosol. Here we showed that the cytosolic and glycosomal pathways are functional. Like in all other eukaryotes, the cytosolic pathways feed glycosylation reactions, however the role of the duplicated glycosomal pathways is currently unknown. We also showed that one of these enzymes (UGP) is imported into glycosomes by piggybacking on another glycosomal enzyme (PEPCK), which are not functionally related. The UGP/PEPCK association is unique since all piggybacking examples reported to date involve functionally related interacting partners, which broadens the possible combinations of carrier-cargo proteins being imported as hetero-oligomers.

## Introduction

*Trypanosoma brucei* is a parasite responsible for human African trypanosomiasis, also known as sleeping sickness, a disease affecting sub-Saharan Africa that can be fatal if left untreated (1). This parasite is transmitted through the bite of a tsetse fly and has a complex developmental cycle including bloodstream (BSF) and procyclic (PCF) forms found in the blood of mammalian hosts and the digestive tract of the insect, respectively. A major difference between these two forms is their mode of energy conservation, with the former depending on glucose *via* glycolysis and the latter being able to use glucose, proline and other amino acids as carbon sources (2). The complexity of *T*. *brucei*’s life cycle leads to a fast and high adaptation capacity to environmental conditions mostly through metabolic changes related to energy metabolism. One of the factors playing a role in these efficient changes is the presence of peroxisome-related organelles called glycosomes. The glycosomes contain the six or seven first glycolytic steps, which are commonly present in the cytosol of other eukaryotic cells (3). In addition, the glycosomes contain up to a dozen of other metabolic pathways, including the sugar nucleotide biosynthetic pathways, which are also exclusively cytosolic in other organisms (4).

All eukaryotes, excepted trypanosomatids, synthesize sugar nucleotides in the cytosol and then transport them into the lumen of the endoplasmic reticulum or Golgi apparatus to feed glycosyltransferase-dependent glycosylation reactions (5). In the particular case of trypanosomatids, most of the enzymes involved in *de novo* biosynthesis of sugar nucleotides are present in the glycosomes (6–11). Some of them are known to be essential for the parasite survival, probably because the cell surface and endosomal/lysosomal systems are rich in essential glycoconjugates (12).

Within the steps involved in production of sugar nucleotides, UDP-glucose pyrophosphorylase (UGP) catalyses the coupling of glucose 1-phosphate (G1P) and UTP to produce UDP-glucose (UDP-Glc) (13). UDP-Glc is a central metabolite that acts as a glucose donor in several pathways, as exemplified by UDP-Glc:glycoprotein glucosyltransferase (UGGT), which uses this sugar nucleotide as a glucosyl donor for protein glycosylation. UDP-Glc has an important role in glycoprotein quality control in the ER, because UGGT specifically glycosylates unfolded glycoproteins to prevent their processing towards the cytosol (14). UDP-Glc is also the obligate precursor of UDP-galactose (UDP-Gal) via a reaction catalysed by UDP-Glc 4’-epimerase (GalE), given that the parasite hexose transporters are unable to transport galactose (15). The lethality of the *T. brucei* GalE null mutant makes UDP-Glc production essential for the parasite (9). In the closely related parasites *T. cruzi* and *Leishmania major*, UDP-sugar pyrophosphorylase (USP) can also activate G1P, in addition to galactose 1-phosphate, while the *T. brucei* genome does not contain the *USP* orthologous gene. Consequently, the simultaneous deletion of the *USP* and *UGP* genes is required to deplete the *Leishmania* cells of UDP-Glc and UDP-Gal, leading to growth arrest and cell death (16). In contrast to the animal and fungal UGP, which are octameric (17) and can be regulated by redox mechanisms (18–20) or phosphorylation (21), the characterized *T. brucei* and *Leishmania major* UGP are active as monomers and are regulated by allosteric mechanisms (7, 17, 22).

The *T. brucei* UGP was reported to be localised in glycosomes of BSF (Marino et al., 2010). However, it does not contain any of the canonical peroxisomal targeting signals (PTS) required for import of proteins into the organelle, *i.e.* the PTS1 tripeptide ([STAGCN]-[RKH]-[LIVMAFY]) or PTS2 ([M]-x_0/20_-[RK]-[LVI]-x_5_-[HQ]-[ILAF], where x refers to any amino acid with their number in subscript) located at the C- and N-terminal extremities of the peroxisomal/glycosomal proteins, respectively (23). Alternatively, proteins lacking a PTS can be imported into the organelle by piggybacking through interaction with a PTS-containing protein. The very few examples of piggybacking described so far in peroxisomes of mammals (24, 25), plants (26) and yeast (27, 28) involve hetero-oligomeric complexes formed by protein isoforms or by functionally related proteins. This mechanism of import has been proposed as an explanation to the presence of some PTS-lacking proteins within glycosomes, but has not yet been reported in trypanosomatids so far.

Here we showed that UGP is imported into glycosomes by interacting with the glycosomal PTS1-containing phosphoenolpyruvate carboxykinase (PEPCK), supporting co-import of functionally unrelated proteins. We also showed that UGP is an essential enzyme for growth of trypanosomes with a dual cytosolic and glycosomal localisation. Metabolomic analyses revealed that UDP-Glc is produced by functional cytosolic and glycosomal pathways. The positive selection of functional sugar nucleotide biosynthesis within glycosomes of trypanosomatids, while this pathway is exclusively cytosolic in the other eukaryotes, raises questions about its role in these parasites.

## Results

### UDP-glucose pyrophosphorylase (UGP) has a dual glycosomal and cytosolic localization

Previous studies on the UGP subcellular localisation revealed that the protein is associated with glycosomes of the BSF (7), despite the absence of any predicted peroxisomal targeting signal (PTS1/PTS2). We raised an anti-UGP immune serum to confirm this unique glycosomal localisation of UGP in PCF by western blotting analyses of glycosomal and cytosolic fractions prepared by differential centrifugation, using as controls antibodies against glycosomal (NADH-dependent fumarate reductase, FRDg) and cytosolic (enolase, ENO) proteins. The anti-UGP immune serum detected a 55 kDa protein corresponding to the predicted size of UGP (theoretical MW: 54.5 kDa), in both glycosomal and cytosolic fractions (Fig. 1A). This dual localisation was further confirmed by digitonin titration as UGP was released together with the cytosolic protein at low concentrations of detergent and the UGP signal increased with the digitonin concentration required to release the glycosomal marker (Fig. 1B). The increased signal at higher digitonin concentrations suggests that the total amount of UGP in the glycosomes is at least equivalent to that in the cytosol. We also addressed the UGP subcellular localisation in BSF by performing hypotonic lysis, which released cytosolic proteins while glycosomal proteins remained in the cellular pellet, as evidenced by the glycosomal aldolase and cytosolic enolase markers (Fig. 1C). UGP is similarly distributed over the two compartments in BSF, as observed for PCF (Fig. 1C).

**Fig 1.**
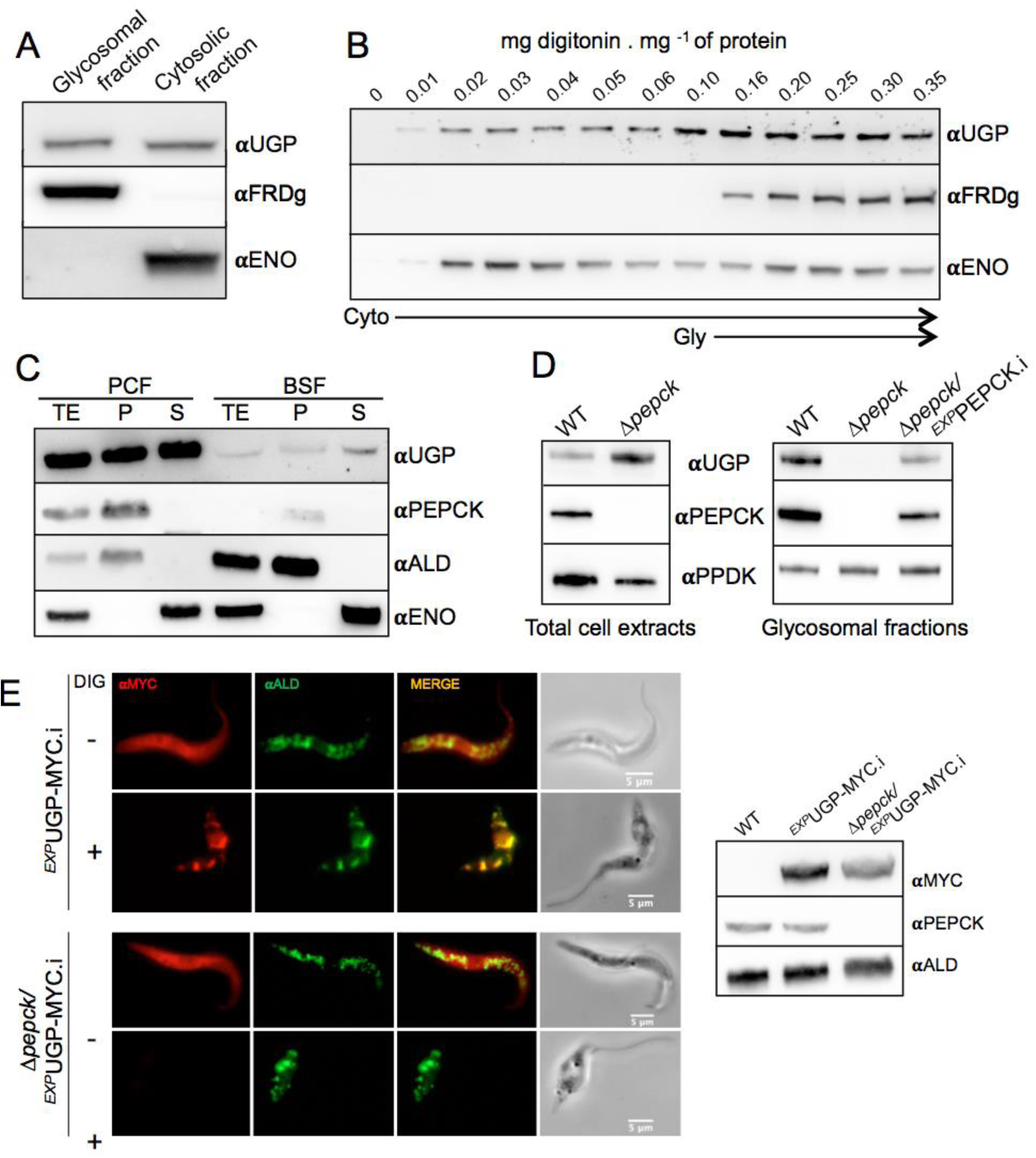
UGP has a dual localisation in PCF and BSF and its import into glycosomes depends on the PTS1-cointaining protein PEPCK. Panels A and B show the subcellular localisation of UGP in the EATRO1125.T7T procyclic trypanosomes. Enriched glycosomal and cytosolic fractions were obtained by differential centrifugation and analysed by western blot (panel A) using the anti-UGP antibodies (αUGP), as well as immune sera against the glycosomal NADH-dependent fumarate reductase (αFRDg) and cytosolic enolase (αENO) markers. The UGP localisation was also studied by digitonin titration (panel B). The supernatants collected from the parental cells incubated with 0-0.35 mg of digitonin per mg of protein were analysed by western blot using the immune sera indicated in the left margin. Panel C compares the subcellular localisation and protein expression levels of UGP and PEPCK, as well as the aldolase glycosomal and enolase cytosolic markers, in PCF and BSF trypanosomes. Total extracts (TE), pellets (P) and supernatants (S) obtained after hypotonic lysis were analysed by western blot using the immune sera indicated. Panel D shows the western blot analysis of total cellular extracts and glycosomal fractions of WT, Δ*pepck* null mutant and the tetracycline-induced Δ*pepck*/*^EXP^*PEPCK rescue cell line (Δ*pepck*/*^EXP^*PEPCK.i) using the anti-UGP (αUGP), anti-PEPCK (αPEPCK) and anti-PPDK (αPPDK, pyruvate phosphate dikinase) immune sera. In the left part panel E, the UGP subcellular localisation was analysed by immunofluorescence of cell lines expressing a recombinant MYC-tagged UGP in the WT (*^EXP^*UGP-MYC cell line) and Δ*pepck* background (Δ*pepck*/*^EXP^*UGP-MYC), using anti-MYC (αMYC, red) and the glycosomal ALD (αALD, green) control. The cells were treated (+ DIG) or not (-DIG) with 0.04 mg digitonin/mg of protein to remove or not the cytosolic UGP-MYC signal, respectively. The expression of UGP-MYC was controlled by western blots on total cell extracts (right panel) using the anti-MYC, anti-PEPCK and anti-ALD as loading control.

### PEPCK-dependent import of UGP into glycosomes

Incidentally, a comparative proteomic analysis of the previously obtained PEPCK null mutant (Δ*pepck*) (29) and the parental cell line, carried out in order to control the *PEPCK* gene deletion, showed a strong reduction (19.7-fold) of UGP peptide counts in the enriched glycosomal fractions of the mutant (see the PXD020190 dataset in the PRIDE partner repository). Depletion of UGP in the glycosomes of the Δ*pepck* cell line was confirmed by western blotting analyses, showing that UGP was no longer detected in the Δ*pepck* glycosomes, while the protein was still present in the total cell extracts (Fig. 1D). Importantly, re-expression of the *PEPCK* gene in the *PEPCK* null background (Δ*pepck*/*^EXP^*PEPCK.i cell line; .i stands for tetracycline-induced) rescued the glycosomal localisation of UGP (Fig. 1D). These data suggest that import of UGP into the glycosomes depends on the presence of PTS1-containing PEPCK, potentially by the so-called piggybacking mechanism not reported so far in trypanosomatids (30). In this context, UGP would be co-transported with PEPCK, which is imported into the glycosome *via* its PTS1.

To confirm this dual subcellular localisation of UGP, we produced cell lines expressing a MYC-tagged UGP under the control of tetracycline in both the parental and the Δ*pepck* backgrounds (Fig. 1E, right panel). Immunofluorescence analyses showed a clear cytosolic pattern in the tetracycline induced *^EXP^*UGP-MYC.i and Δ*pepck*/*^EXP^*UGP-MYC.i cell lines (Fig. 1E, left panel). A signal co-localising with the glycosomal marker aldolase was detected for the *^EXP^*UGP-MYC.i cells only after treatment with 0.04 mg of digitonin per mg of protein required for permeabilization of the plasma membrane. These data confirmed that the recombinant UGP-MYC exhibits a dual localisation similar to the native protein. Interestingly, the glycosomal signal was not detected in the Δ*pepck*/UGP-MYC.i cell line after digitonin treatment, indicating that all UGP localises exclusively in the cytosol of this mutant. Altogether, these data support the role of PEPCK in the import of UGP into glycosomes.

### UGP interacts with PEPCK in some glycosomes

To evidence the putative interaction between UGP and PEPCK we used Proximity Ligation Assays (PLA Duolink), which enables detection of protein interactions including transient/weak interactions *in situ* with high specificity and sensitivity (31). We produced a Δ*pepck*/*^EXP^*TY-PEPCK/*^EXP^*UGP-MYC cell line expressing the TY-tagged PEPCK (TY-PEPCK) and MYC-tagged UGP (UGP-MYC) in the *PEPCK* null background. Briefly, the second *PEPCK* allele of the single allele Δ*pepck/*PEPCK knockout cell line was replaced by a *TY-PEPCK* copy encoding TY-PEPCK tagged at its N-terminal extremity to preserve the PTS1 motif required for glycosomal import. This Δ*pepck*/*^EXP^*TY-PEPCK cell line was transfected with the pLew100- *^EXP^*UGP-MYC plasmid to express UGP-MYC under the control of tetracycline. As controls, the UGP-MYC and TY-PEPCK recombinant proteins have been independently expressed in the Δ*pepck* (Δ*pepck*/*^EXP^*UGP-MYC) and parental (*^EXP^*TY-PEPCK) backgrounds, respectively. The expression of both recombinant proteins, the specificity of the primary antibodies and the glycosomal import of TY-PEPCK and UGP-MYC were confirmed by western blot analyses of enriched glycosomal and cytosolic fractions, digitonin titration and immunofluorescence analyses (Fig. S1). As expected, TY-PEPCK showed a glycosomal localisation, however its level of expression was ∼8 times lower than that of the native protein (Fig. S1A, compare the upper and lower band of αPEPCK signal in the *^EXP^*PEPCK-TY cell line, respectively). Despite this difference in expression levels, a significant part of the recombinant UGP-MYC is imported into the glycosomes of the tetracycline induced Δ*pepck*/*^EXP^*TY-PEPCK/*^EXP^*UGP-MYC.i cell line (Fig. S1A,B), while remaining exclusively in the cytosol of the Δ*pepck*/*^EXP^*UGP-MYC.i cell line (Fig. S1A), as previously shown (Fig. 1E). αMYC (Rabbit) and αTY (mouse) were validated to be specific and sensitive enough to perform PLA analysis (Fig. S1C).

PLA positive puncta (red signal) corresponding to TY-PEPCK/UGP-MYC hetero-oligomers were observed in 62% of the Δ*pepck*/*^EXP^*TY-PEPCK/*^EXP^*UGP-MYC.i cells, while only 7% and 6% of the control Δ*pepck*/*^EXP^*UGP-MYC.i and *^EXP^*PEPCK-TY.i cells were positive, respectively (Fig. 2A-B). In addition, ∼90% of the positive Δ*pepck*/*^EXP^*UGP-MYC.i and *^EXP^*PEPCK-TY.i cells contained a single dot and the other 10% contained 2 dots, while the number of dots per cell in the Δ*pepck*/*^EXP^*TY-PEPCK/*^EXP^*UGP-MYC.i population was much higher, with 62% of the cells showing 2 to 10 dots (Fig. 2B). These data are in agreement with interactions between UGP-MYC and TY-PEPCK in Δ*pepck*/*^EXP^*TY-PEPCK/*^EXP^*UGP-MYC.i cells, while the very few red dots observed within the control Δ*pepck*/*^EXP^*UGP-MYC.i and *^EXP^*PEPCK-TY.i cells represent background signals. Staining with an immune serum against the glycosomal PPDK showed that the PLA signals are found very close to the PPDK-containing organelles without showing clear co-localisation with them (Fig. 2C). This suggests the existence of different pools of glycosomes, as previously reported (32).

**Fig 2.**
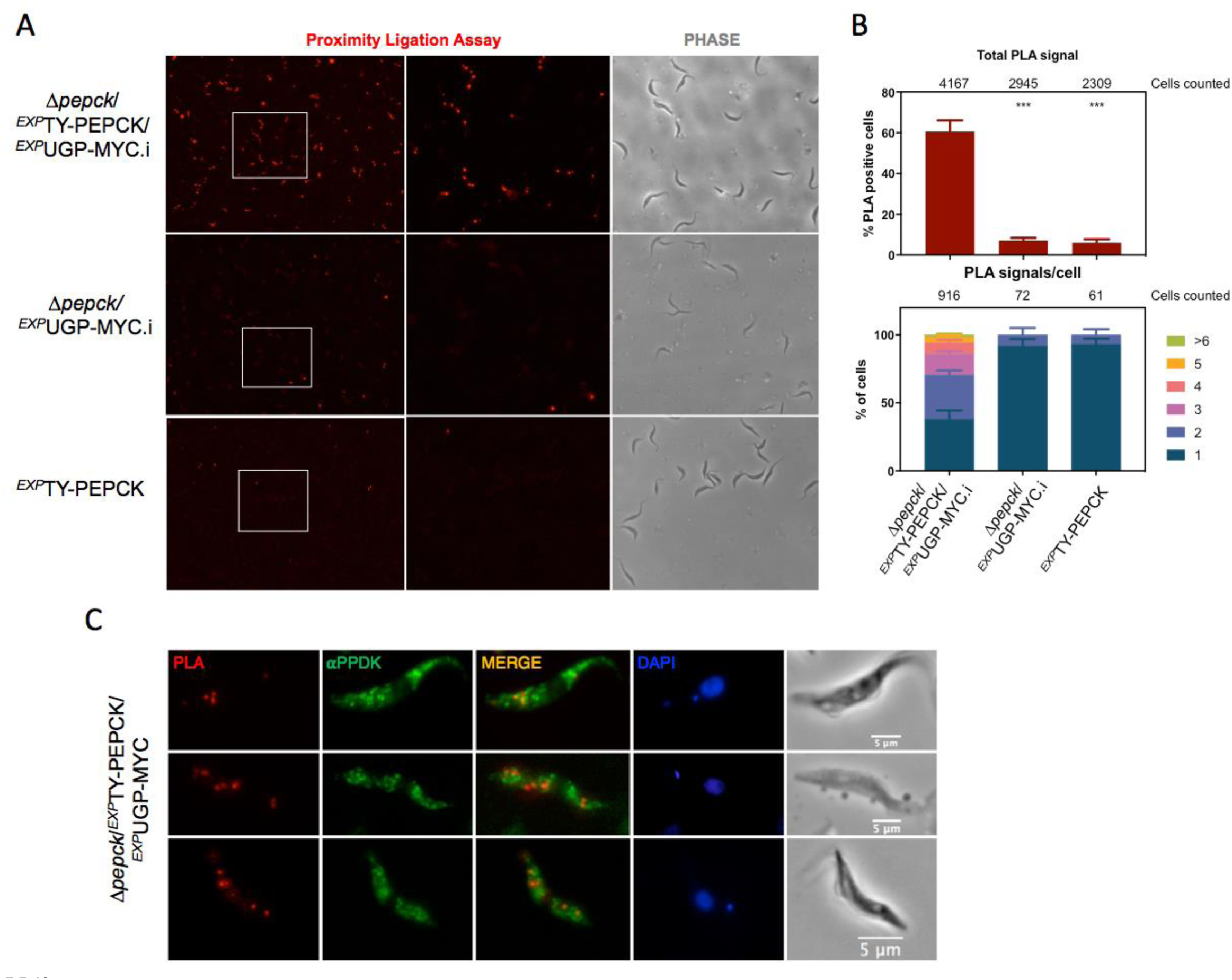
UGP interacts transiently with PEPCK. Panel A shows *in situ* proximity ligation assay (PLA) analysis of the interaction between MYC-tagged UGP and TY-tagged PEPCK in the Δ*pepck*/*^EXP^*TY-PEPCK/*^EXP^*UGP-MYC.i cell line. The *^EXP^*PEPCK-TY and Δ*pepck*/ *^EXP^*UGP-MYC.i cell lines expressing only one recombinant protein were used as negative controls. The central (PLA) and right (phase contrast) panels are enlargements of the white rectangles shown in the left panel. In panel B, the % of PLA positive cells is shown for each cell line and the total cell number counted is indicated on the top of the graph. The percentage values correspond to the average of 12 pictures randomly taken from 2 independent experiments. Significant differences between samples are indicated: ***p < 0.001. The number of PLA signals per cell was analysed by counting manually the number of dots using ImageJ for positive cells (lower panel). In panel C, the localisation of PLA signal was analysed in detail and compared with the PPDK glycosomal marker (αPPDK) by counterstaining after the Duolink *In Situ* Protocol.

### Determination of critical parts for PEPCK-UGP interaction

To investigate which part of UGP and PEPCK interacts with its piggybacking partner, truncated versions of each protein were expressed in the parental or Δ*pepck* cell lines, respectively. Since PEPCK form homodimers (33), the truncated PEPCK proteins were expressed in the Δ*pepck* cell line to prevent heterodimer formation. UGP is reported to be monomeric (7, 22) and was only detected as monomer in native gel analyses (Fig. S2), therefore the native and recombinant proteins will not directly interact. We expressed in the parental background the recombinant UGP with the 10xTY tag either at the N-terminal or the C-terminal ends of UGP (*^EXP^*TY-UGP_1-485_ and *^EXP^*UGP_1-485_-TY cell lines, respectively), by *in situ* replacement of one *UGP* allele. The subcellular distribution of UGP in these cell lines was determined by western blot analyses of glycosomal and cytosolic fractions. The N-terminal tag affected glycosomal import of UGP, since the glycosomal localisation of TY-UGP_1-485_ was decreased by ∼9-fold compared to the native UGP in parental cells (Fig. 3A, left panel). However, no changes were observed in the glycosome:cytosol ratio for the UGP_1-485_-TY (Fig. 3A, right panel). C-terminus tagged recombinant UGP versions truncated from their N-terminal (UGP_xxx-485_-TY, Fig. 3B) or C-terminal (UGP_1-XXX_-TY, Fig. 3C) extremities were inserted *in situ* to produce new cell lines. It is to note that the UGP coding sequence used for the UGP_XXX-485_-TY constructs was recoded from position 165 to 337 amino acids to become resistant to the RNAi construct (see below), which was useful to confirm the correct insertion of the recombinant fragment in the *UGP* locus (Fig. S3). The UGP_124-485_-TY truncated protein was no more imported inside the glycosomes (Fig. 3B), while glycosomal import of the UGP_1-124_-TY, UGP_1-173_-TY and UGP_1-226_-TY proteins was not affected (Fig. 3C), suggesting that the N-terminal domain up to the 123 amino acid position contains residues interacting with PEPCK. The truncated recombinant UGP missing (UGP_66-485_-TY) or containing only (UGP_1-66_-TY) the N-terminal 66 residues were imported into glycosomes, although with a lower efficiency compared to the parental cell line, suggesting that key residues of the PEPCK-binding site are located on either side of position 66 (Fig. 3B-C). The presence of the PEPCK binding site in the N-terminal extremity of UGP may explain the low glycosomal import of the TY-UGP_1-485_ recombinant protein (Fig. 3A).

**Fig 3.**
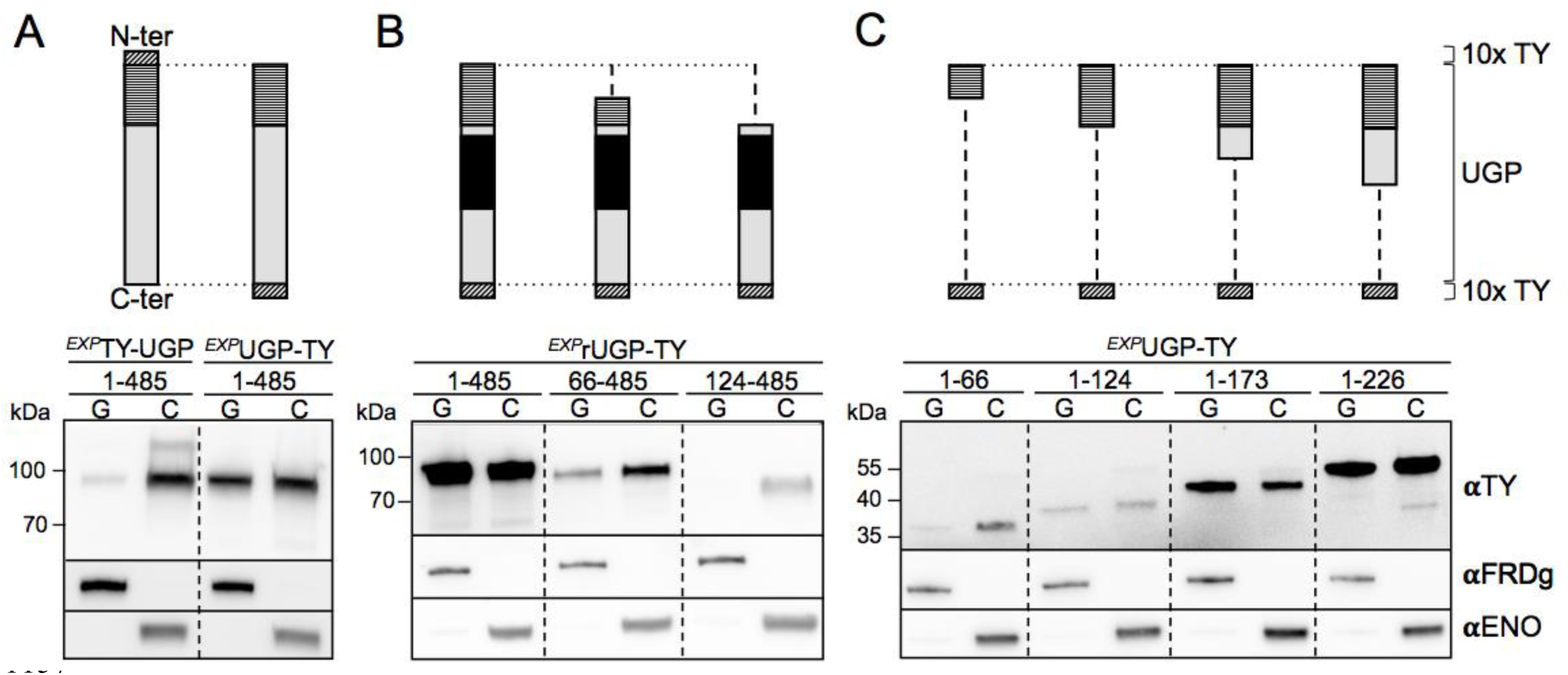
The N-terminal 123 residues of UGP are required for import into the glycosomes. The top of the figure shows schematic representations of the 10xTY-tagged native or recoded UGP and the corresponding truncations. The 123-residues peptide required for UGP import, the recoded part of UGP and the 10xTY tags are highlighted by horizontally hatched boxes, obliquely hatched boxes and black boxes, respectively. The lower part of the figure shows western blot analyses of glycosomal (G) and cytosolic (C) fractions produced from cell lines expressing 10xTY-tagged recombinant UGP, using the anti-TY antibody (αTY) as well as immune sera from glycosomal (αFRDg) and cytosolic (αENO) markers. Panel A shows UGP recombinant proteins tagged at the N-terminal (*^EXP^*TY-UGP_1-485_) or C-terminal (*^EXP^*UGP_1-485_-TY) extremities, while panels B and C show truncated UGP tagged at the C-terminal extremity. The truncations designed from the N-terminal end lack the first 66 (66-485) or 123 (124-485) residues, while truncations designed from the C-terminal end contain the N-terminal first 66 (1-66), 124 (1-124), 173 (1-173) or 226 (1-226) residues (Panel B). A PCR analysis was performed to confirm the correct insertion of the UGP_XXX-485_-TY fragments at the *UGP* locus (Fig. S3)

We performed a similar analysis to determine the PEPCK region involved in UGP glycosomal import by expressing truncated versions of recombinant PEPCK using the pLew100 vector. The PEPCK was truncated from its N-terminal extremity in order to maintain the C-terminal PTS1 required for glycosomal import of both PEPCK and UGP. Unfortunately, none of the truncated PEPCK peptides were detectable by western blotting in total cell extracts, probably due to protein instability. To resolve this stability issue, the truncated PEPCK peptides were fused to the C-terminal extremity of the green fluorescent protein (eGFP) and used to produce four different cell lines (Fig. 4A). We determined the glycosomal import of UGP in these Δ*pepck*/*^EXP^*eGFP-PEPCK_XXX-525_ cell lines by western blot analyses of glycosomal and cytosolic fractions. As mentioned above, UGP is no more detected in glycosomes isolated from the parental Δ*pepck* mutants (Fig. 4B). The glycosomal import of UGP was not affected in the absence of the N-terminal first 140 and 180 residues of PEPCK (Δ*pepck*/*^EXP^*eGFP-PEPCK_140-525_ and Δ*pepck*/*^EXP^*eGFP-PEPCK_180-525_ cell lines), while deletion of the N-terminal first 214 and 321 residues abolished glycosomal import of UGP, which remained exclusively in the cytosolic fractions (Fig. 4C). This suggested that the 34-residues peptide between amino acids positions 180 and 214 of PEPCK is required for UGP import into glycosomes. Importantly, none of the eGFP-PEPCK truncations have PEPCK activity, indicating that the import of UGP is not related to PEPCK activity inside the glycosomes (Fig. 4D)

**Fig. 4.**
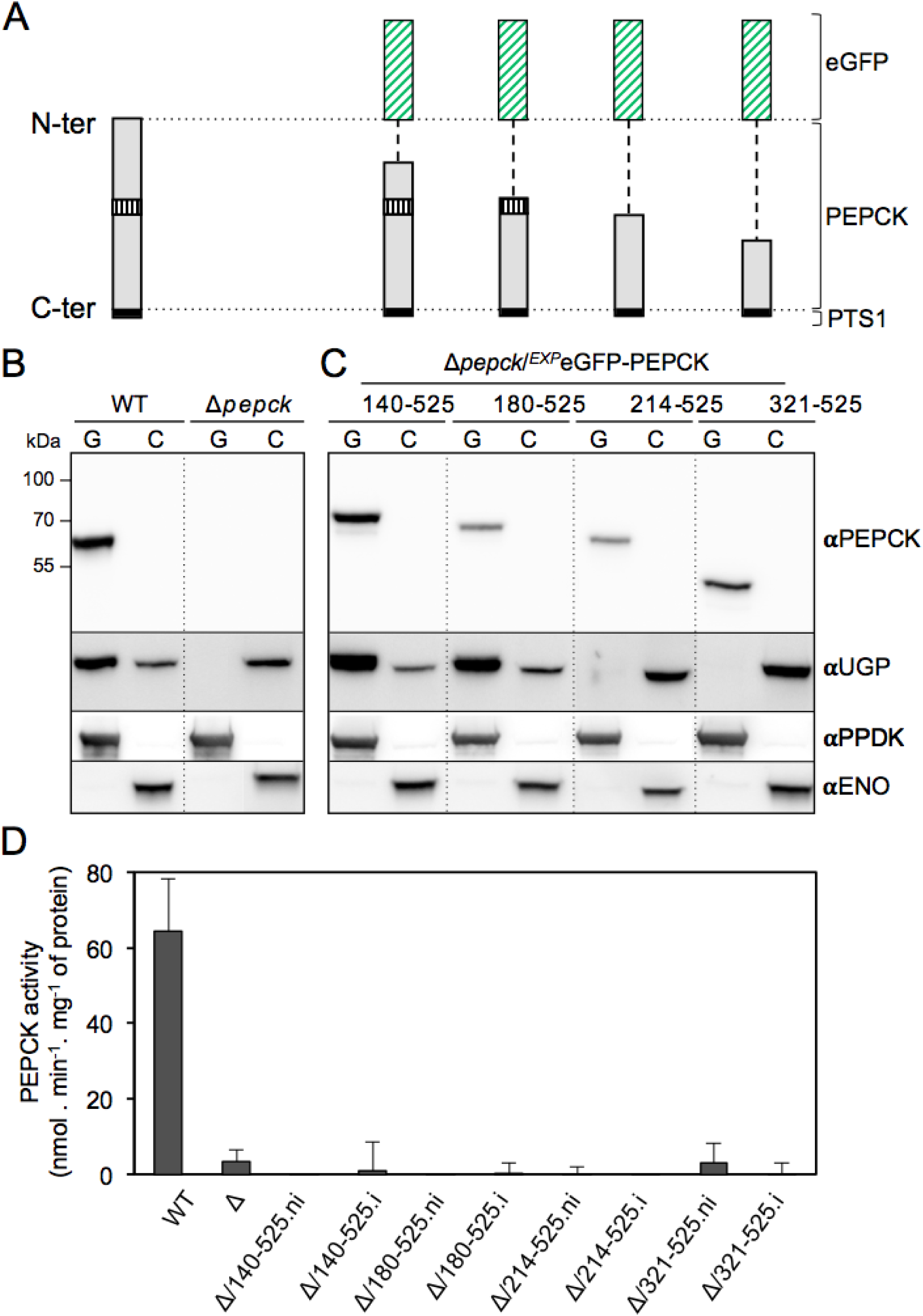
A 34-residues peptide of PEPCK is required for glycosomal import of UGP. Panel A shows schematic representations of the endogenous PEPCK and eGFP-PEPCK recombinant proteins expressed in the Δ*pepck* background, in which the 34-residues peptide required for UGP import (vertically hatched boxes) is highlighted. Panels B and C show western blot analyses of glycosomal (G) and cytosolic (C) fractions obtained from the parental and Δ*pepck* cell lines (panel B), as well as from cell lines expressing truncated eGFP-PEPCK recombinant proteins (Δ*pepck*/*^EXP^*eGFP-PEPCK_XXX-525_), with anti-PEPCK (αPEPCK) and anti-UGP (αUGP) immune sera. Glycosomal (αPPDK) and cytosolic (αENO) markers are used to check the quality of glycosomal and cytosolic fractions. Panel D shows PEPCK activity determined in total extracts of WT, Δ*pepck* (Δ) and the non-induced (.ni) and induced (.i) Δ*pepck*/ *^EXP^*eGFP-PEPCK_XXX-525_ cell lines.

### The UGP protein is essential for *T. brucei*

The stem-loop RNAi strategy was used with the conditional pLew100 vector to address the role of UGP in the procyclic trypanosomes. Two *^RNAi^*UGP cell lines obtained from individual transfections (H10 and E4) showed a strong reduction of growth 7 days after tetracycline induction, indicating that UGP is essential for PCF viability (Fig. 5A, top panel). For both RNAi cell lines, the growth rate of the parental strain was restored 18 days post-induction, concomitantly with the re-expression of the native UGP (Fig. 5A, lower panel). This re-expression of RNAi targeted genes is often observed for trypanosome essential genes (29). It is noteworthy that the UGP expression was barely detectable in the non-induced *^RNAi^*UGP-H10 total cell extracts. Western blotting analyses of enriched glycosomal fractions, which proved to be more sensitive than on total cell extracts, showed that UGP expression was reduced by ∼30-fold compared to the parental cells without any significant effect on growth (Fig. 5B, left panel). This suggests that UGP activity is present in large excess in parental PCF. The distribution of UGP between glycosomal and cytosolic compartments is not affected by this ∼30-fold reduction (Fig. 5B). After five days of induction, UGP was no more detectable in the glycosomal fractions and was reduced by ∼2-fold in the cytosol (Fig. 5B). These small amounts of UGP were not sufficient to sustain growth of PCF.

**Fig. 5.**
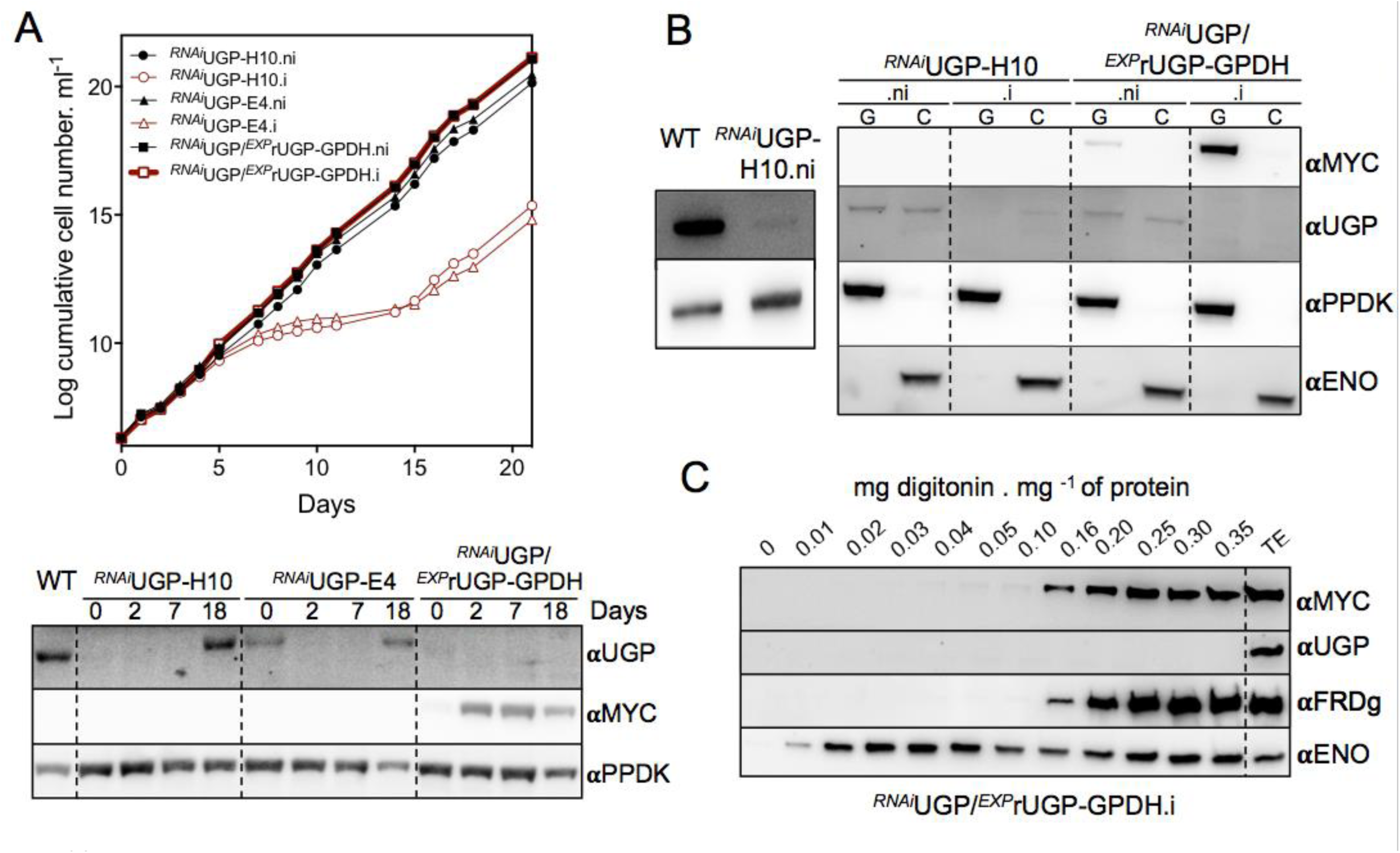
Production and functional analyses of *^RNAi^*UGP cell lines. Panel A shows a growth curve of the tetracycline-induced (.i) and non-induced (.ni) *^RNAi^*UGP-H10, *^RNAi^*UGP-E4 and *^RNAi^*UGP/*^EXP^*rUGP-GPDH cell lines. The expression of the native UGP and recombinant rUGP-GPDH upon induction was monitored by western blot analysis using anti-UGP (αUGP) and anti-MYC (αMYC) immune sera, respectively, and anti-PPDK (αPPDK) as loading control (lower panel). Panel B shows western blot analyses of glycosomal (G) and cytosolic (C) fractions produced from the parental cells (WT), as well as tetracycline-induced (.i) and non-induced (.ni) *^RNA^*^i^UGP-H10 and *^RNAi^*UGP/*^EXP^*rUGP-GPDH cell lines using immune sera described in precedent figures. The left panel was used to quantify relative expression of UGP in glycosomes of non-induced *^RNA^*^i^UGP-H10 mutant. The glycosomal localisation of recombinant rUGP-GPDH (αMYC) was confirmed by western blot analyses of supernatants obtained after digitonin titration of *^RNAi^*UGP/*^EXP^*rUGP-GPDH.i cell line, using immune sera described in precedent figures (panel C). The control lane TE corresponds to total extract from *^EXP^*rUGP-GPDH.i cell line.

To determine whether UGP is also required for growth of the procyclic trypanosomes in the insect-like glucose-free conditions, the parasites were grown in the absence of glucose, as described before (34). The growth of the *^RNAi^*UGP.i and Δ*pepck*/*^RNAi^*UGP.i cell lines is similar regardless of the amounts of glucose in the medium (Fig. S4), indicating the UGP is probably also essential in the insect vector, which is considered to be free of glucose (35). In addition, the subcellular distribution of UGP in the parental cells is not affected by the absence of glucose (Fig. S5).

### Targeting a recombinant UGP exclusively to the glycosomes

To elucidate in which subcellular compartment the UDP-Glc/UDP-Gal biosynthetic pathway is active (glycosomes and/or cytosol), it was necessary to express UGP exclusively in the cytosol or in the glycosomes of the parasite. The exclusive cytosolic localisation of UGP in the viable Δ*pepck* mutant demonstrated that UGP is functionally active in the cytosol. To assess the role of UGP in glycosomes, we optimized the glycosomal import of UGP and expressed it in the *^RNAi^*UGP background. To do so, a recombinant *UGP* gene recoded to become resistant to the RNAi construct (*rUGP*) was fused at its 3’-extremity with a 3xMYC tag followed by different glycosomal targeting peptides (PTS1), namely the last C-terminal 12 residues of glycosomal FRDg (*rUGP-FRDgPTS1*), the full-length PTS1-containing glycosomal glycerol-3-phosphate dehydrogenase (*GPDH*) gene (*rUGP-GPDH*), and the full-length PTS1-containing glycosomal phosphoglycerate (*PGKc*) gene (*rUGP-PGKc\**). Since glycosomal expression of PGK is lethal for the PCF trypanosomes (36), the codon of the lysine residue (K215) essential for the PGK enzymatic activity (37) was replaced by the alanine codon. These recombinant proteins were conditionally expressed in the parental cell line and their distribution between the glycosomal and cytosolic compartments was determined by digitonin titration (Fig. 6A). The rUGP-FRDgPTS1 and cytosolic enolase proteins showed the same cytosolic profiles, which implies that the extended FRDg PST1 motif is not sufficient for glycosomal import of UGP. In contrast, the rUGP-PGKc* recombinant protein is mostly associated to the glycosomes, but a minor part remained in the cytosol. Finally, the rUGP-GPDH (∼100 kDa) and the glycosomal FRDg proteins were both released with a minimum of 0.16 mg digitonin per mg protein, which is consistent with the exclusively glycosomal localisation of this recombinant protein. The UGP activity was increased by 5-fold in the *^EXP^*rUGP-GPDH.i cell line compared to the non-induced (.ni) and parental cell lines, which validated the functionality of the rUGP-GPDH protein (Fig. 6B). Expression of the rUGP-GPDH had no effect on morphology, growth and survival of the *^EXP^*rUGP-GPDH cell line. Consequently, rUGP-GPDH can be expressed in the *^RNAi^*UGP background to address the role of UGP in the glycosomes.

**Fig. 6.**
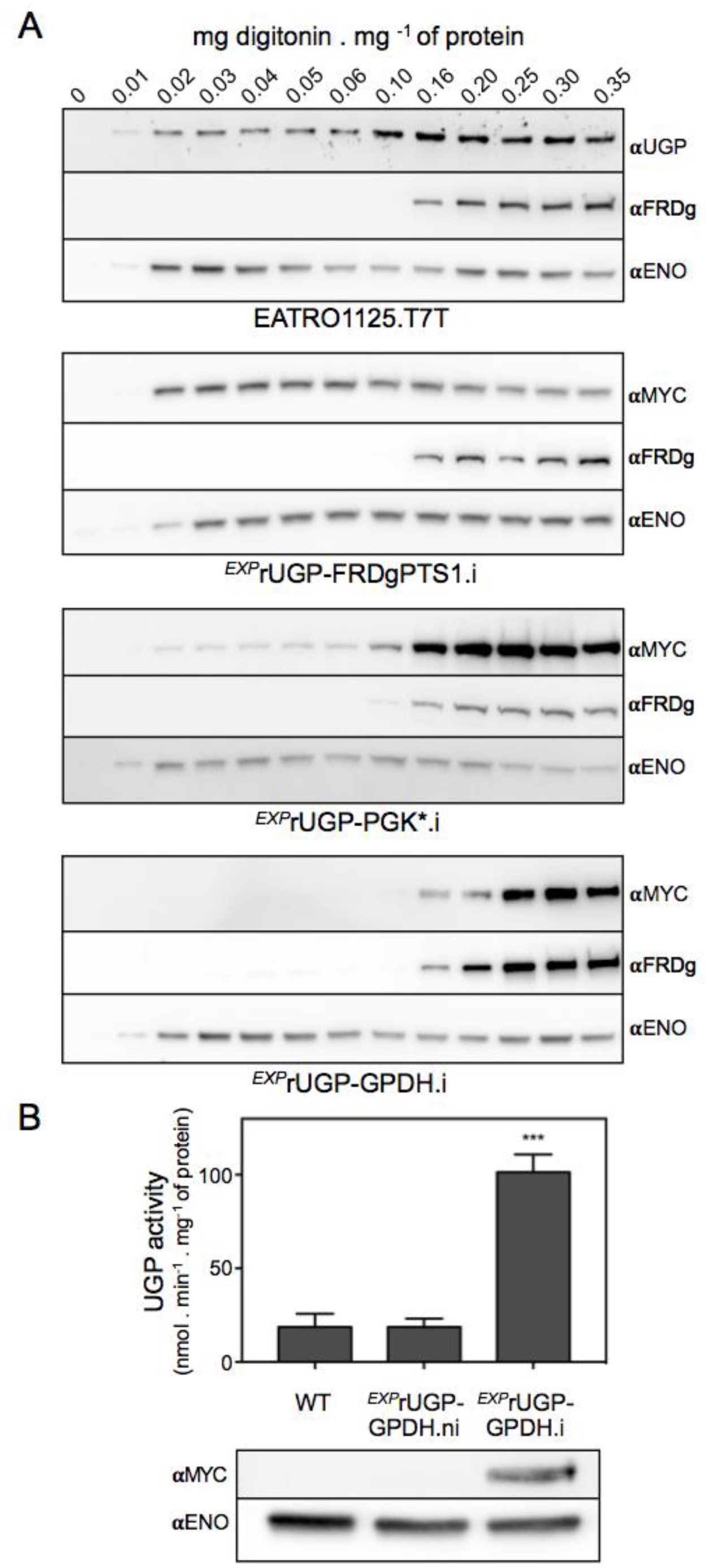
Expression of a glycosomal recombinant UGP. In panel A, the subcellular localisation of the recombinant UGP was monitored by western blot analysis of supernatants obtained after digitonin titration of the tetracycline induced (.i) *^EXP^*rUGP-FRDgPTS1, *^EXP^*rUGP-PGKc* and *^EXP^*rUGP-GPDH cell lines using anti-MYC (αMYC) antibody. The anti-UGP immune serum (αUGP) was used to detect UGP in parental cells fractions (top panel). Anti-FRDg (αFRDg) and anti-ENO (αENO) immune sera were used as glycosomal and cytosolic markers, respectively. Panel B shows the UGP activity measured in total cell extracts of parental (WT) and tetracycline-induced (.i) and non-induced (.ni) *^EXP^*rUGP-GPDH cell lines (n=3, SEM). Significant differences between samples are indicated: ***p < 0.001. Western blot analysis of *^EXP^*rUGP-GPDH expression with anti-MYC (αMYC) and anti-ENO (αENO, loading control) immune sera is shown below the graph.

### Expression of the glycosomal rUGP-GPDH rescues the lethality of the *^RNAi^*UGP mutant

The *^EXP^*rUGP-GPDH construct (pHD1336-rUGP-GPDH), which produces an exclusively glycosomal rUGP, was introduced into the *^RNAi^*UGP-H10 cell line. Western blot analyses showed that native UGP was no longer detectable in glycosomal and cytosolic fractions of *^RNAi^*UGP/*^EXP^*rUGP-GPDH.i cell line, while the dying *^RNAi^*UGP.i cells still expressed residual amounts of UGP in the cytosol (Fig. 5B, right panel). The exclusive glycosomal subcellular localisation of the recombinant rUGP-GPDH protein in the *^RNAi^*UGP/*^EXP^*rUGP-GPDH.i cell line observed by cellular fractionation (Fig. 5B) was confirmed by digitonin titration (Fig. 5C). In this context of absence of cytosolic UGP, the viability of the *^RNAi^*UGP/*^EXP^*rUGP-GPDH.i cell line (Fig. 5A) strongly supports the hypothesis that the glycosomal pathway is functional.

### The Δ*ugp*/*^EXP^*rUGP-GPDH cell line is viable

Considering that UGP is an essential protein, knock-out mutants were produced in two cell lines expressing tetracycline-inducible recombinant UGP, *i.e.* glycosomal/cytosolic rUGP (recoded UGP followed by a MYC tag) and glycosomal rUGP-GPDH. The *UGP* alleles were replaced by the PAC and BLE markers after transfection with the recombinant plasmids expressing rUGP (Δ*ugp*/*^EXP^*rUGP) or rUGP-GPDH (Δ*ugp*/*^EXP^*rUGP-GPDH), in the presence of tetracycline to express the recombinant rUGP or rUGP-GPDH, respectively. Deletion of both *UGP* alleles was confirmed by PCR (Fig. 7A) and western blot (Fig. 7B) analyses. Tetracycline removal did not induce death of the parasites (Fig. 7C), since the recombinant rUGP and rUGP-GPDH proteins were still expressed after 18 days in the absence of tetracycline (Fig. 7C inset and Fig. 7B). However, the growth of the Δ*ugp*/*^EXP^*rUGP-GPDH cell line is slightly affected after tetracycline removal, which is consistent with the essential role of UGP. More importantly, the viability of the double mutant Δ*ugp*/*^EXP^*rUGP-GPDH supports our hypothesis that the glycosomal pathway is functional.

**Fig. 7.**
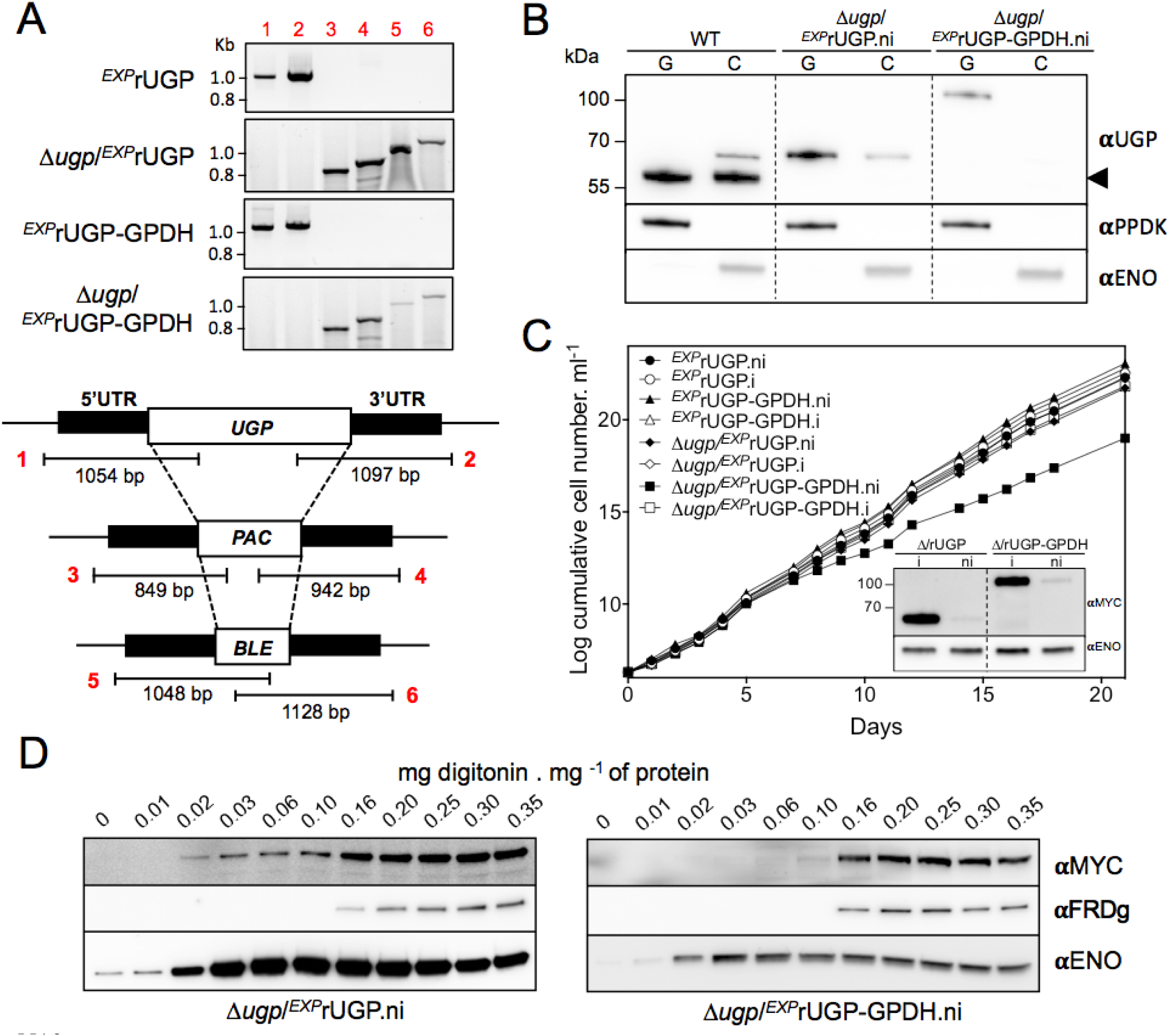
Production and functional analyses of Δ*ugp* cell lines. Panel A shows a PCR analysis of genomic DNA isolated from the parental (*^EXP^*rUGP and *^EXP^*rUGP-GPDH) and null mutant (Δ*ugp*/*^EXP^*rUGP and Δ*ugp*/*^EXP^*rUGP-GPDH) cell lines. Both knock-outs were obtained in the presence of tetracycline. Primers are designed on sequences flanking the 5’UTR and 3’UTR fragments used to target *UGP* gene depletion (black boxes) and on the ORF of the *UGP* gene, as well as the puromycin (*PAC*) and phleomycin (*BLE*) resistance genes (white boxes). The Δ*ugp*/*^EXP^*rUGP represents a control cell line expressing a recombinant UGP with a dual cytosolic and glycosomal localisation. Panel B shows glycosomal (G) and cytosolic (C) fractions obtained after subcellular fractionation of the UGP null cell lines in the absence of tetracycline (5 days). The arrowhead highlights the native UGP only in parental (WT) cells. In panel C, the growth of the cell lines was followed during 21 days, in the presence (.i) or the absence (.ni) of tetracycline. Western blot analyses with anti-MYC (αMYC) and anti-ENO (αENO, loading control) of the tetracycline-induced (.i) and non-induced (.ni), 18 days after tetracycline removal) Δ*ugp*/*^EXP^*rUGP and Δ*ugp*/*^EXP^*rUGP-GPDH mutants are shown in the inset. In panel D, the UGP null cell lines were analysed by digitonin titration 5 days after removal of tetracycline. Western blot of supernatants confirmed the exclusive glycosomal localisation of recombinant rUGP-GPDH with anti-MYC (αMYC) in Δ*ugp*/*^EXP^*rUGP-GPDH cell line and the dual localisation of rUGP in Δ*ugp*/*^EXP^*rUGP cell line.

These data in agreement with the functional role of the glycosomal UGP, have to be confirmed by determining the subcellular localisation of rUGP-GPDH in the Δ*ugp*/*^EXP^*rUGP-GPDH cell line. After 5 days in the absence of tetracycline, the viable Δ*ugp*/*^EXP^*rUGP-GPDH.ni cell line expresses the recombinant rUGP-GPDH exclusively in the glycosomes (Fig. 7B-D, right panels). These data confirmed that the UDP-Glc/UDP-Gal biosynthetic pathway, which includes UGP, is active in the glycosomes. As expected, the MYC-tag rUGP showed a dual glycosomal and cytosolic localisation in the Δ*ugp*/*^EXP^*rUGP cell line (Fig. 7B-D).

### The glycosomal and cytosolic UGP-containing pathways are functional

To confirm the functionality of the glycosomal and cytosolic pathways involving UGP, cell lines expressing the native and/or recombinant UGP in both subcellular compartments (WT, *^EXP^*rUGP.i and *^EXP^*rUGP-GPDH.i), only in the cytosol (Δ*pepck*), only in the glycosomes (*^RNAi^*UGP/*^EXP^*rUGP-GPDH.i) or not at all (*^RNAi^*UGP-H10.i) were further analysed (Fig. 8A-C). This included determining the expression levels of UGP in the glycosomal and cytosolic fractions by western blotting and enzymatic activities, as well as quantifying intracellular metabolites, including the substrate (G1P) and the product (UDP-Glc) of the UGP enzymatic reaction, by mass-spectrometry based metabolomics profiling approach (IC-HRMS).

**Fig 8.**
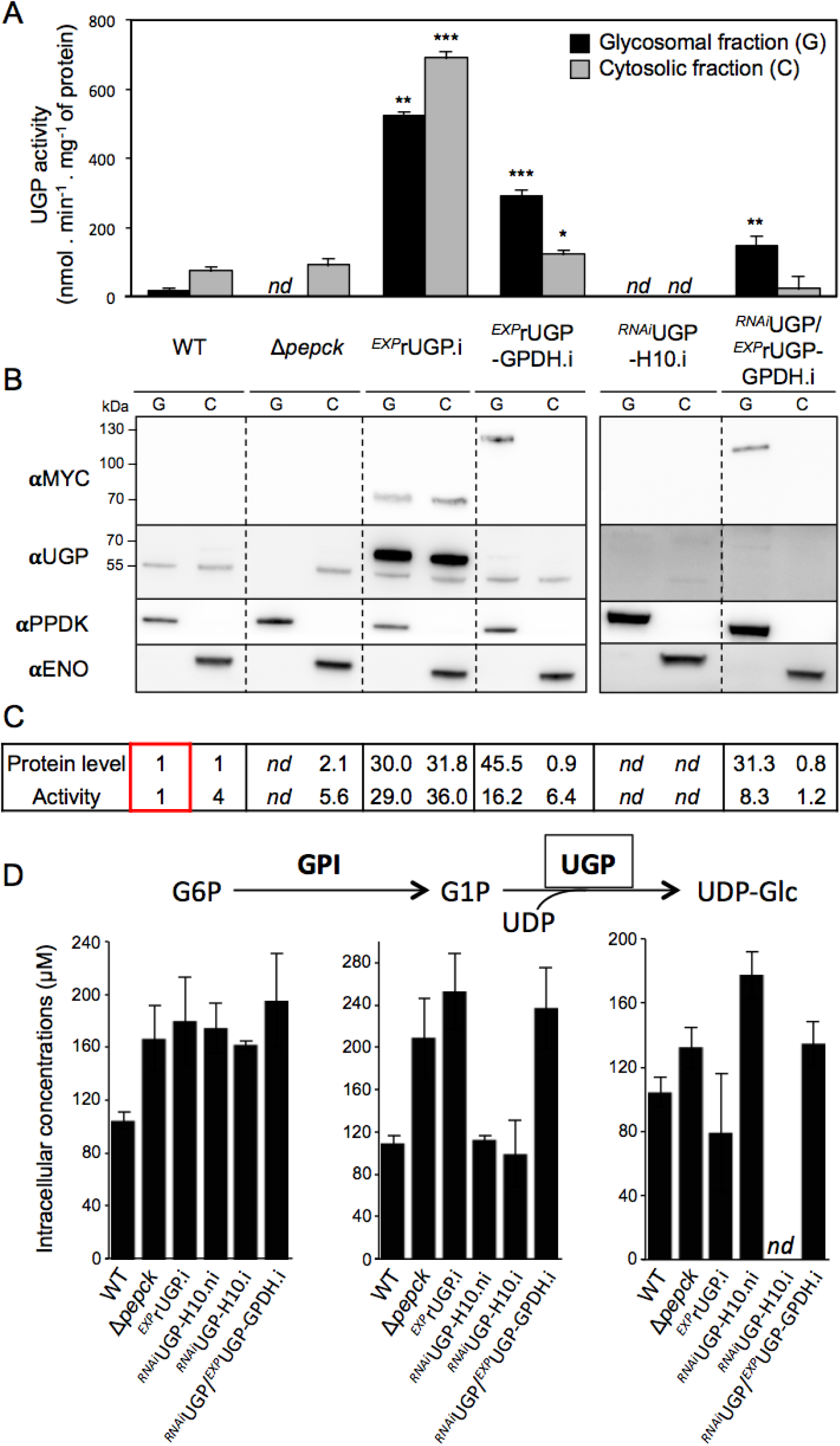
PCF produces UDP-Glc in glycosomes and the cytosol. UGP activity was determined in enriched glycosomal and cytosolic fractions of the WT, Δ*pepck, ^EXP^*rUGP.i, *^EXP^*rUGP-GPDH.i, *^RNAi^*UGP-H10.i and *^RNAi^*UGP/*^EXP^*rUGP-GPDH.i cell lines (Panel A, n=3, SEM). The cytosolic and glycosomal UGP activities were normalized to the cytosolic malic enzyme and the glycosomal glycerol kinase activities, respectively. Significant differences between WT and mutants are indicated for each compartment: ***p < 0.001, **p < 0.01, *p < 0.05. Panel B shows a representative western blot analysis of the corresponding cell lines. The recombinant rUGP-GPDH was detected with anti-MYC antibody (αMYC). Glycosomal (αPPDK) and cytosolic (αENO) markers are also shown. Panel C shows relative amounts of UGP determined by western blot and specific activity (average of n=3). For these comparative analyses, the protein and activities levels detected in the cytosolic fraction of the WT cells were used as references with the arbitrary value of 1 (boxed values). Panel D shows IC-HRMS analyses of intracellular metabolites (G6P, G1P and UDP-Glc) collected from indicated cell lines incubated in glucose-rich SDM79 medium. Only G6P, G1P and UDP-Glc are shown in this figure, for other hexose phosphates and triose phosphates see Figure S6. *nd*, non-detectable; .i, tetracycline induced; .ni, non-induced.

The specific activity of UGP (ratio between enzymatic activity and relative amount of proteins detected by western blotting) in the cytosolic fractions of *^EXP^*rUGP.i is ∼3.5-times lower than in the parental WT cells, suggesting that the C-terminal MYC tag affects UGP activity (Fig. 8C). Similarly, the native UGP shows a specific activity in the glycosomal fraction 4-times lower than in the cytosolic fraction of the parental cells, which suggests that the glycosomal sequestration of UGP affects its activity by a yet unknown mechanism. We also confirmed that the coupling enzyme (UDP-Glc dehydrogenase) used in the UGP activity assays was not affected by the presence of the same amounts of the glycosomal or cytosolic samples (273 *versus* 245 mU.mg^-1^ of protein, respectively). The activity of the recombinant rUGPs, which is ∼30 times more expressed in the *^EXP^*rUGP.i than the native UGP, was not affected in glycosomes, as the enzyme specific activity is similar in the glycosomal and the cytosolic fractions (Fig. 8C). It is also noteworthy that the UGP activity was detected in the cytosol of the *^RNAi^*UGP/*^EXP^*rUGP-GPDH.i, while the native UGP is not detectable by western blot (Fig. 8B) and the recombinant rUGP-GPDH is exclusively glycosomal (Fig. 5C). This could be due to the rupture of a few glycosomes during the grinding step designed to primarily disrupt the plasma membrane.

To confirm the role of UGP subcellular localisation in UDP-Glc production, we used mass-spectrometry based metabolomics to determine the intracellular amounts of G6P, G1P and UDP-Glc (Fig. 8D), as well as other metabolites as controls (Fig. S6) in the cell lines mentioned above cultivated in SDM79 medium. This metabolomics approach was validated with the analysis of the Δ*pepck* cell line, in which the metabolic flux through the Gly3P/DHAP shuttle, used to maintain the glycosomal redox balance, has been reported to be increased in the absence of PEPCK (29). Indeed, the level of Gly3P is increased by ∼3-times in the Δ*pepck* mutant compared to all the other cell lines analysed (Fig. S6). Regarding the sugar nucleotide biosynthetic pathways, only UDP-Glc and UDP-GlcNAc were identified and quantified with this methodology (Fig. S6) and the levels of UDP-Glc detected (80-170 uM) are comparable to those previously reported for procyclic trypanosomes (110 to 540 uM) (38) (Fig. 8D). UDP-Glc is no longer detectable in the *^RNAi^*UGP.i cell line (Fig. 8D), which shows that UGP is the only enzyme producing UDP-Glc in PCF trypanosomes. It is also of note that UDP-Glc is detected in non-induced *^RNAi^*UGP cells at levels similar to parental cells, despite the ∼30-fold reduction of UGP protein levels (Fig. 6B), which shows that PCF trypanosomes express a large excess of UGP. Most importantly, UDP-Glc is produced in cells expressing UGP exclusively in the cytosol (Δ*pepck*) or in glycosomes (*^RNAi^*UGP/*^EXP^*rUGP-GPDH.i) at levels similar to those of WT cells, which confirms the functionality of the pathway in both subcellular compartments.

## Discussion

Trypanosomatids are known to sequester a cascade of consecutive glycolytic enzymes into glycosomes, in addition to enzymes of other pathways including gluconeogenesis, the pentose phosphate pathway and sugar nucleotide biosynthesis (4, 39). In this study, we address three questions related to the glycosome biology by analysing UGP, a key enzyme of the sugar nucleotide biosynthesis involved in UDP-Glc synthesis: (*i*) the physiological role of this glycosomal pathway remains unknown since it is also present in the cytosol, the subcellular compartment where biosynthesis of sugar nucleotides takes place in the other eukaryotes, (*ii*) the molecular mechanisms leading to the import of glycosomal enzymes lacking peroxisomal targeting signals (PTS1 or PTS2) have not been investigated so far in trypanosomatids and (*iii*) mammalian peroxisomes multiply by the ER *de novo* route or by growth and division followed by protein import into newly produced organelles, but what about glycosomes? Here we show that (*i*) the glycosomal pathway leading to production of UDP-Glc and UDP-Gal is functional and is essential in the PCF trypanosomes, in the absence of the cytosolic pathway, (*ii*) UGP is imported into glycosomes by piggybacking on the PTS1-containing PEPCK and (*iii*) PEPCK and UGP interact in only a few glycosomes, which may represent newly produced glycosomes competent for protein import.

### What is the role of sugar-nucleotide biosynthesis in glycosomes?

The functionality of the glycosomal and cytosolic UGP-containing pathways was validated by the viability of mutants expressing UGP exclusively in glycosomes (*^RNAi^*UGP/*^EXP^*rUGP-GPDH.i) or cytosol (Δ*pepck*) and the detection of UDP-Glc in both cell lines. This first direct evidence of a functional production of sugar nucleotides inside glycosomes raises two questions. First, how the *de novo* synthesized UDP-Glc and UDP-Gal leave the glycosomes to reach the ER and Golgi apparatus where they are required for protein glycosylation? The glycosomal membrane is considered to be impermeable to bulky metabolites such as nucleotides, since the size limitation of the general peroxisomal diffusion pore is in the order of 400 Da (39, 40). Consequently, exchange of sugar nucleotides between the glycosomal and cytosolic compartments requires transporters. However, the only transporters known to be associated with the glycosomal membrane are the ABC transporters GAT1, GAT2 and GAT3, with GAT1 likely transporting acyl-CoAs (41, 42) and proteomics analyses of glycosomal membrane fractions did not reveal additional candidates (43). Further work is certainly required to confirm the presence of such sugar nucleotide transporters in the glycosomal membrane. Second, what is the role of sugar-nucleotide biosynthesis inside the glycosomes, since the cytosolic pathway is functional in the procyclic trypanosomes, as observed in all eukaryotes? UGP has also been localised in the Golgi apparatus, chloroplasts, membrane fractions, as well as in the cell walls where it also provides UDP-Glc to produce glycoconjugates in plants and yeasts (44). Interestingly, the yeast UGP also shows a dual subcellular localisation depending on phosphorylation at the N-terminus S11 residue, with the non-phosphorylated cytosolic and phosphorylated cell wall enzymes being involved in glycogenesis and cell wall glucan synthesis, respectively (21). All of these biosynthetic pathways require glycosyltransferases, which have not been detected in the glycosomal proteomes (45, 46) or in the repertoire of PTS-containing proteins (47). This supports the view that UDP-Glc and UDP-Gal are not produced in the glycosomes to feed glycosylation reactions inside glycosomes. Alternatively, glycosomal UDP-Glc could have a signalling role, as previously observed in animals and plants (48, 49).

### Piggybacking is a low efficient import process, as observed for UGP

Piggybacking has been described as an import mechanism with relatively low efficiency in four out of five examples of physiological hetero-oligomer import into peroxisomes reported so far, *i.e.* superoxide dismutase (SOD1) (24) and lactate dehydrogenase (LDH) (25) in mammals, pyrazinamidase/nicotinamidase (PNC1) (28) and malate dehydrogenase 2 (Mdh2) (27) in yeast, which are co-imported with the PTS-containing copper chaperone SOD1 (CCS), read-through-extended LDH (LDHBx), glycerol-3-phosphate dehydrogenase (GPD1) and Mdh3, respectively. These four co-imported proteins display dual peroxisomal and cytosolic localisation with the majority remaining within the cytosol (27, 50). Similarly, approximately half of UGP remains in the cytosol. The reason of this relatively low import efficiency has been elucidated by the demonstration that the PST1 receptor (PEX5), required for peroxisomal import of PTS1-containing proteins, binds preferentially to monomers compared to oligomers (51). Interestingly, weak protein-protein interactions are sufficient to support piggyback import. Indeed, blue native gels failed to show interaction between the mammalian SOD and CCS partners (24) and synthetic substrates designed to evaluate the import of proteins showed dissociation constants (K_d_) differing over three orders of magnitude, with even an apparent K_d_ of ∼6×10^−3^ M allowing the detection of piggyback import (52). Despite several attempts, we did not observe any interaction between UGP and PEPCK using co-immunoprecipitation or native gels, suggesting that these interactions are weak and transient. In agreement with this weak interaction, PEPCK is in large excess compared to UGP, as illustrated by the ∼30-fold higher enzymatic activity of PEPCK compared to UGP (670 *versus* 20 mU.mg^-1^ of protein) (53) and the ∼100-fold higher peptide counts for PEPCK than UGP in proteomics analyses of glycosomal fractions from PCF (see the PXD020190 dataset in the PRIDE partner repository). In conclusion, our results support the role of hetero-oligomer import by piggybacking as an alternative route for import of glycosomal proteins, as described for peroxisomes of mammals and yeast. More importantly, the UGP/PEPCK association provides the first example of hetero-oligomeric import by piggybacking involving two proteins not functionally related, since PEPCK is involved in the maintenance of the glycosomal redox and ATP/ADP balances, as well as gluconeogenesis (29, 36). Indeed, among the other known examples of piggybacking, CCS is the chaperone of SOD1 (24), LDH and LDHBx are encoded by the same gene (25), Mdh2 and Mdh3 are Mdh isoforms (27), the PST1-containing phosphatase B subunit and phosphatases A/C subunits form an heterotrimeric enzymatic complex (26), however, the peroxisomal functions of PNC1 and GPD1 are unknown (28).

### UGP and PEPCK would interact only transiently upon their import into newly produced import competent glycosomes

Since the formation of the UGP/PEPCK heterodimer may occur mainly during UGP import into the organelle, the analysis of the UGP/PEPCK interactions using the PLA approach provides new insights regarding glycosomal import of proteins and multiplication of the organelles. In mammalian cells, peroxisomes multiply by the ER *de novo* route and by growth and division. This latter case involves an asymmetric process generating new peroxisomes *via* formation of a membrane compartment and subsequent import of newly synthesised matrix proteins (54–56). Indeed, overexpression of the membrane peroxin Pex11pβ resulted in the formation in mammalian cells of pre-peroxisomal membrane structures composed of mature globular domains and tubular extensions, the latter being maturated by import of matrix proteins (55). Equivalent clusters of tubular glycosomal membranes were also observed by overexpressing Pex11 in *T. brucei* (57) and clusters of elongated glycosomes have more recently been observed in BSF trypanosomes by whole cell reconstruction using 3D electron microscopy (58). In addition, *T. brucei* expresses Fis1 and Dpl1, two key proteins involved in the fission of newly produced peroxisomes in other eukaryotes (59–61). Overall, these observations confirm that glycosomes multiply by growth and division as observed for the mammalian peroxisomes. This also implies that the new peroxisomes/glycosomes produced by growth and division are the most competent organelles for protein import and that they represent only a limited fraction of the organelle population, supporting the heterogeneity observed before among the peroxisomal (62) and glycosomal (32) populations. We thus propose that the structures showing UGP/PEPCK close proximity by PLA correspond to newly produced import-competent glycosomes. Considering that (*i*) PEPCK and UGP mainly physically interact during import at the glycosomal membrane, because of their weak and transient interaction, (*ii*) that only up to 10 dots per cell correspond to physical proximity between PEPCK and UGP, with most cells containing 2-5 dots (Fig. 2), while PEPCK and UGP appear localised in almost all, if not all, glycosomes (Fig. 1E and Fig. S1) and (*iii*) that the number of glycosomes was estimated to 60-65 per G1 trypanosome cell (58, 63), one could consider that the 3-10% of the organelles showing UGP/PEPCK interaction by PLA are newly produced glycosomes importing the matrix proteins, including PEPCK and UGP, in this context.

## Materials and Methods

### Trypanosomes and cell cultures

The procyclic form of *T. brucei* EATRO1125.T7T (TetR-HYG T7RNAPOL-NEO) was cultured at 27°C in SDM79 medium containing 10% (v/v) heat inactivated fetal calf serum, 5 μg.ml^-1^ hemin (64), hygromycin (25 µg.ml^-1^) and neomycin (10 µg.ml^-1^). Alternatively, the cells were cultivated in a glucose-free medium derived from SDM79, called SDM79-GlcFree (34). The bloodstream form of *T. brucei* 427 90–13 (TetR-HYG T7RNAPOL-NEO) was cultured at 37°C in IMDM supplemented with 10% (v/v) heat-inactivated fetal calf serum (FCS), 0.25 mM ß-mercaptoethanol, 36 mM NaHCO_3_, 1 mM hypoxanthine, 0.16 mM thymidine, 1 mM sodium pyruvate, 0.05 mM bathocuprone and 2 mM L-cysteine (65). Cells were transfected as previously described (Bringaud et al., 1998). Over-expression and RNAi cell lines were induced with tetracycline (1 µg.ml^-1^). Growth was followed by daily cell counting with the cytometer Guava EasyCyte.

### Expression of MYC-tagged UGP, TY-tagged UGP, eGFP-PEPCK truncations and TY-tagged PEPCK

The *UGP* gene (Tb927.10.13130) was cloned using the In-Fusion cloning system (Clontech) in the HindIII-NdeI restriction sites of pLew100-X-MYC, which was designed for expression of recombinant protein tagged at the C-terminal extremity with 3 MYC epitopes (modified from (66)). The EATRO1125.T7T parental cell line, Δ*pepck* (29), TY-PEPCK and Δ*pepck*/TY-PEPCK cell lines were transfected with the pLew100-UGP-MYC tetracycline inducible plasmid and cells were selected in SDM79 containing phleomycin (5 μg.ml^-1^). The *UGP* gene was also *in situ* tagged at the N-terminal or C-terminal extremities (67). Briefly, the DNA sequence encoding 10xTY1 tag and BLA resistance cassette was amplified from the pPOTv7-10xTY1 vector using long primers (see Table S1) that incorporate a 5’ overhang of 80 nt homologous to *UGP* gene and its UTR. For production of UGP truncated versions 10xTY1 tagged at their C-terminal extremity, the forward primers were designed within the *UGP* gene extension to produce proteins containing the N-terminal first 66 (1-66), 124 (1-124), 173 (1-173) and 226 (1-226) residues. The PCR products were precipitated with ethanol before use for transfection and cells were selected in SDM79 containing blasticidin (20 μg.ml^-1^). We also expressed truncated versions of a recoded UGP (rUGP; see Fig. S7) lacking either the first 66 or 124 residues fused to 10xTY1 tag at their C-terminal extremity. The PCR fragments corresponding to complete or truncated *rUGP* gene fused to TY tag and blasticidin cassette from pPOTv7 were obtained by overlapping PCR and cloned into pGEM-T. Cells were transfected with 10 ug of plasmid digested with NotI. For expression of eGFP-PEPCK truncated versions, the Δ*pepck* cell line (29) was transfected with the pLew100 tetracycline inducible plasmid containing truncated versions of PEPCK fused at the N-terminal extremity with eGFP to increase the stability of the recombinant truncated proteins. PCR fragments corresponding to the truncations of PEPCK at residues 140, 180, 214 and 321 were inserted between the XhoI and XbaI restriction sites of the pLew100-eGFPX plasmid using the In-Fusion cloning system (Clontech). The PEPCK gene was also *in situ* tagged at the N-terminal as described above.

### Production of glycosomal recombinant UGP proteins

To target UGP exclusively to the glycosomes, the recoded recombinant *UGP* (*rUGP*; see Fig. S7) gene was inserted in the pHD1336 expression vector (41). For this purpose, the rUGP was fused at its C-terminal extremity to 3xMYC tag followed by (*i*) the sequence encoding the last C-terminal 12 residues of the glycosomal fumarate reductase (*FRDg*) gene, which contains a PTS1 (rUGP-FRDgPST1), (*ii*) the full-length PTS1-containing glycosomal phosphoglycerate (*PGKc*) gene (rUGP-PGKc*) and (*iii*) the full-length PTS1-containing glycerol-3-phosphate dehydrogenase (*GPDH*) gene (rUGP-GPDH). The K215 residue, essential for PGK activity (37), was replaced by alanine. In order to increase the net charge of residues at the C-terminal, which is a major determinant of peroxisomal import efficiency (68), we modified one residue in the C-terminal extremity of PGK (TL**R**NRW-SSL instead of TL**S**NRW-SSL) and of GPDH (PA**R**PRT-SKM instead of PA**L**PRT-SKM). The pHD1336-rUGP-FRDgPST1 plasmid, provided by the GeneCust Company, was used for cloning the synthesized genes (GeneCust) *PGKc\** and *GPDH* in the MluI-BamHI restriction sites. The EATRO1125.T7T parental cell line was transfected and cells were selected in SDM79 containing blasticidin (20 μg.ml^-1^).

### Inhibition of *UGP* gene expression

The inhibition of UGP expression by RNAi was achieved by expression of stem-loop “sense/antisense” RNA molecules targeting a 537-bp fragment of *UGP* gene introduced in the pLew100 tetracycline inducible expression vector. A PCR-amplified 579-bp fragment, containing the antisense UGP sequence was inserted between HindIII and BamHI restriction sites of pLew100 plasmid. Then, the separate 537-bp PCR-amplified fragment containing the sense *UGP* sequence was inserted upstream the antisense sequence, using HindIII and XhoI restriction sites. The resulting plasmid pLew-UGP-SAS contains a sense and antisense version of the UGP fragment separated by a 42-bp fragment. The *^RNAi^*UGP and *^RNAi^*UGP/*^EXP^*rUGP-GPDH mutants were generated by transfecting the EATRO1125.T7T and *^RNAi^*UGP cell lines with the pLew-UGP-SAS plasmid and the pHD1336-rUGP-GPDH plasmid, respectively. Transfected cells were selected in SDM79 medium containing hygromycin (25 µg.ml^-1^), neomycin (10 µg.ml^-1^) and phleomycin (5 µg.ml^-1^), with addition of blasticidin (20 μg.ml^-1^) for the *^RNAi^*UGP/*^EXP^*rUGP-GPDH cell line.

### Production of UGP null mutants

Replacement of the *UGP* gene by the phleomycin and puromycin resistance markers via homologous recombination was performed with DNA fragments containing the resistance marker gene flanked by the UTR sequences. Briefly, an HpaI DNA fragment containing the *PAC* or *BLE* resistance marker gene preceded by the UGP 5’UTR fragment (522 bp) and followed by the UGP 3’UTR fragment (526 bp) was cloned in the pGEM-T plasmid. The UGP knock-out mutants were generated in the *^EXP^*rUGP-GPDH and *^EXP^*rUGP cell lines in the presence of tetracycline. The *^EXP^*rUGP cell line was generated by transfecting the EATRO1125.T7T parental cell line with the pHD1336 vector expressing the *rUGP* sequence followed by a MYC tag sequence under the control of tetracycline. Transfected cells were selected in SDM79 medium containing blasticidin (20 µg.ml^-1^), phleomycin (5 µg.ml^-1^), puromycin (1 µg.ml^-1^) and tetracycline (1 µg.ml^-1^). The selected cell lines *rUGP::GPDH^Ti^-BLA TetR-HYG T7RNAPOL-NEO* Δ*ugp*::*PAC/*Δ*ugp*::*BLE* and *rUGP^Ti^-BLA TetR-HYG T7RNAPOL-NEO* Δ*ugp*::*PAC/*Δ*ugp*::*BLE* are called Δ*ugp/ ^EXP^*rUGP and Δ*ugp*/*^EXP^*rUGP-GPDH, respectively.

### Preparation of glycosomal and cytosolic fractions

Cell homogenates were obtained by grinding pre-washed cells with silicon carbide (200 mesh) in STE buffer (25 mM Tris, 1 mM EDTA, 250 mM sucrose, pH 7.8) (69) supplemented with the Complete EDTA-Free protease inhibitor cocktail (Roche). The cells were microscopically checked for at least 90% disruption. The lysates were diluted in 7 ml of STE, centrifuged at 1,000 g and then at 5,000 g for 10 min each at 4°C. The supernatants were centrifuged at 42,000 g for 10 min at 4°C to yield the glycosome-enriched pellets and the cytosolic fractions (supernatants). The glycosomal pellets were washed once with 1 ml of STE before centrifugation at 42,000 g for 10 min at 4°C and resuspension in 0.2 ml of STE. Equivalent amounts of protein from glycosomal (G) and cytosolic (C) fractions were analysed by western blotting.

### Digitonin permeabilization

Trypanosomes were washed two times in cold PBS and resuspended at 10 mg of protein per ml in STE buffer supplemented with 150 mM NaCl and protease inhibitors. Cell aliquots (100 μl) were incubated with increasing quantities of digitonin (Sigma) for 4 min at 25°C, before centrifugation at 14,000 g for 2 min. The supernatants were analysed by western blotting.

### Cell fractionation by hypotonic lysis

BSF and PCF parasites (2.10^8^) were washed in PBS and hypotonically lysed in presence of protease inhibitors by incubating in 5 mM Na_2_HPO_4_, 0.3 mM KH_2_PO_4_ for 30 min at 4°C before centrifugation at 14,000 g for 15 min. The pellet was solubilized in SDS 2% and both supernatant and pellet were analysed by western blotting.

### Immunofluorescence and Proximity Ligation Assay (PLA)

Cells were washed twice with PBS and then treated (+) or not (-) with 0.04 mg of digitonin per mg of protein during 4 min at 25°C. After centrifugation at 14,000 g for 2 min and washing, the cellular pellets were resuspended in PBS and fixed with 4% paraformaldehyde (PFA) for 10 min at RT. The cells were spread on poly-L-lysine-coated slides and permeabilized with 0.05% Triton X-100. After incubation in PBS containing 4% BSA overnight, cells were incubated for 45 min with primary antibodies (Table S2), washed with PBS and incubated for 45 min with secondary antibodies (Table S2). Slides were washed and mounted with SlowFade Gold (Molecular Probes). Images were acquired with Metamorph software on a Zeiss Imager Z1 or Axioplan 2 microscope as previously described (70).

*In situ* proximity ligation assay (PLA) was performed using the Duolink™ *In Situ* Red Starter Kit Mouse/Rabbit (Sigma-Aldrich) following the manufacturer’s recommendations. Briefly, PFA-fixed and Triton X-100 permeabilized cells were spread on slides as described above. The cells were blocked with Duolink Blocking Solution for 60 min at 37°C. Primary antibodies rabbit anti-MYC (1/1000) and mouse anti-TY1 (1/5000) were diluted in Duolink Antibody Diluent and incubated 60 min at room temperature. The slides were washed for 10 min in wash buffer A and incubated with PLUS and MINUS PLA probes for 60 min at 37°C. Ligation and amplification steps were performed according to the manufacturer’s instructions. After washing, cells were blocked with PBS-BSA 4% overnight. The cells where counterstained with mouse anti-PPDK (αPPDK) (71). Slides were mounted in Duolink *in Situ* Mounting Medium with DAPI. Images were acquired as described for immunofluorescence and analysed using ImageJ. Cells were counted manually using cell counter ImageJ plugin.

### Blue-native PAGE (BN-PAGE)

10^8^ cells were washed in PBS and resuspended in SoTE (0.6 M sorbitol, 2 mM EDTA, 20 mM Tris-HCl, pH 7.5) (72). Cells were incubated with 0 or 0.16 mg of digitonin per mg of protein for 4 min at 25°C, before centrifugation at 14,000 g for 2 min. The supernatants containing both the cytosolic and glycosomal proteins were analysed by BN-PAGE on a precast (3–12%) Bis-Tris polyacrylamide gel (Invitrogen) according to standard methods.

### Western blot analyses

Total protein extracts (5 x 10^6^ cells), glycosomal and cytosolic fractions, or supernatants obtained after digitonin treatment were separated by SDS-PAGE (10%) and immunoblotted on TransBlot Turbo Midi-size PVFD Membranes (Bio-Rad) (73). Immunodetection was performed as described (73, 74) using the primary antibodies and conditions summarized in Table S2. Revelation was performed using the Clarity western ECL Substrate as described by the manufacturer (Bio-Rad). Images were acquired and analysed with the ImageQuant LAS 4000 luminescent image analyser.

### UGP activity assay

The UGP activity in total lysates and aliquots of glycosomal and cytosolic fractions was measured as previously described (22). For normalisation of the UGP activities, the malic enzyme activity was determined on the total cell extracts and the cytosolic fractions, as described before (75). For normalisation of the UGP activities in glycosomal extracts, the glycerol kinase activity was determined as described before (76). The PEPCK activity was measured in total lysates as previously described (Hunt and Köhler, 1995).

### Label-free quantitative proteomics

Enriched glycosomal fractions were loaded on a 10% acrylamide SDS-PAGE gel and proteins were visualized by Colloidal Blue staining. The steps of sample preparation, protein digestion and LC-MS parameters used for nanoLC-MS/MS analysis on a Q-Exactive were previously described (77). For protein identification, Sequest HT and Mascot 2.4 algorithms through Proteome Discoverer 1.4 Software (Thermo Fisher Scientific Inc.) were used for protein identification in batch mode by searching against a *Trypanosoma brucei* protein database (11 119 entries, release 46). This database was downloaded from http://tritrypdb.org website. Two missed enzyme cleavages were allowed. Mass tolerances in MS and MS/MS were set to 10 ppm and 0.02 Da. Oxidation of methionine, acetylation of lysine and deamidation of asparagine and glutamine were searched as dynamic modifications. Carbamidomethylation on cysteine was searched as static modification. Peptide validation was performed using Percolator algorithm (78) and only “high confidence” peptides were retained corresponding to a 1% False Discovery Rate (FDR) at peptide level. Raw LC-MS/MS data were imported in Progenesis QI (version 2.0; Nonlinear Dynamics, a Waters Company) for feature detection, alignment, and quantification. All sample features were aligned according to retention times by manually inserting up to fifty landmarks followed by automatic alignment to maximally overlay all the two-dimensional (m/z and retention time) feature maps. Singly charged ions and ions with higher charge states than six were excluded from analysis. All remaining features were used to calculate a normalization factor for each sample that corrects for experimental variation. Peptide identifications (with FDR<1%) were imported into Progenesis. Only non-conflicting features and unique peptides were considered for calculation of quantification at protein level. A minimum of two peptides matched to a protein was used as the criteria for identification as a differentially expressed protein. The mass spectrometry proteomics data have been deposited to the ProteomeXchange Consortium (http://proteomecentral.proteomexchange.org) via the PRIDE partner repository (79) with the dataset identifier PXD020190.

### Mass spectrometry analyses of intracellular metabolites by IC-HRMS

Parental and mutant cell lines grown in SDM79 medium were collected on filters by fast filtration preparation (2 x 10^7^ cells per filter), as described before (29). Metabolites were analysed by liquid anion exchange chromatography Dionex ICS-5000+ Reagent-Free HPIC (Thermo Fisher Scientific, Sunnyvale, CA, USA) system coupled to Thermo Scientific LTQ Orbitrap Velos hybrid FT mass spectrometer (Thermo Fisher Scientific, San Jose, CA, USA). The metabolites were separated within 48 min, using linear gradient elution of KOH applied to an IonPac AS11 column (250 x 2 mm, Dionex) equipped with an AG11 guard column (50 x 2 mm, Dionex) at a flow rate of 0.35 ml.min^-1^. The column and autosampler temperature were 30°C and 4°C respectively. Injected sample volume was 15 µl. Mass detection was carried out in a negative electrospray ionization (ESI) mode. The settings of the mass spectrometer were as follows: spray voltage 2.7 kV, capillary and desolvatation temperature were 350 and 350 °C respectively, maximum injection time 50 ms. Nitrogen was used as sheath gas (pressure 50 units) and auxiliary gas (pressure 5 units). The automatic gain control (AGC) was set at 1e6 for full scan mode with a mass resolution of 60,000 (at 400 *m/z*). Data acquisition was performed using Thermo Scientific Xcalibur software. The identification of metabolites relied upon matching accurate masses from FTMS scan (mass tolerance of 5 ppm) and retention time using TraceFinder 3.2 software. The absolute levels of intracellular metabolites were quantified based on isotope-dilution mass spectrometry (IDMS) approach.

## Supplementary data

**Fig. S1.**
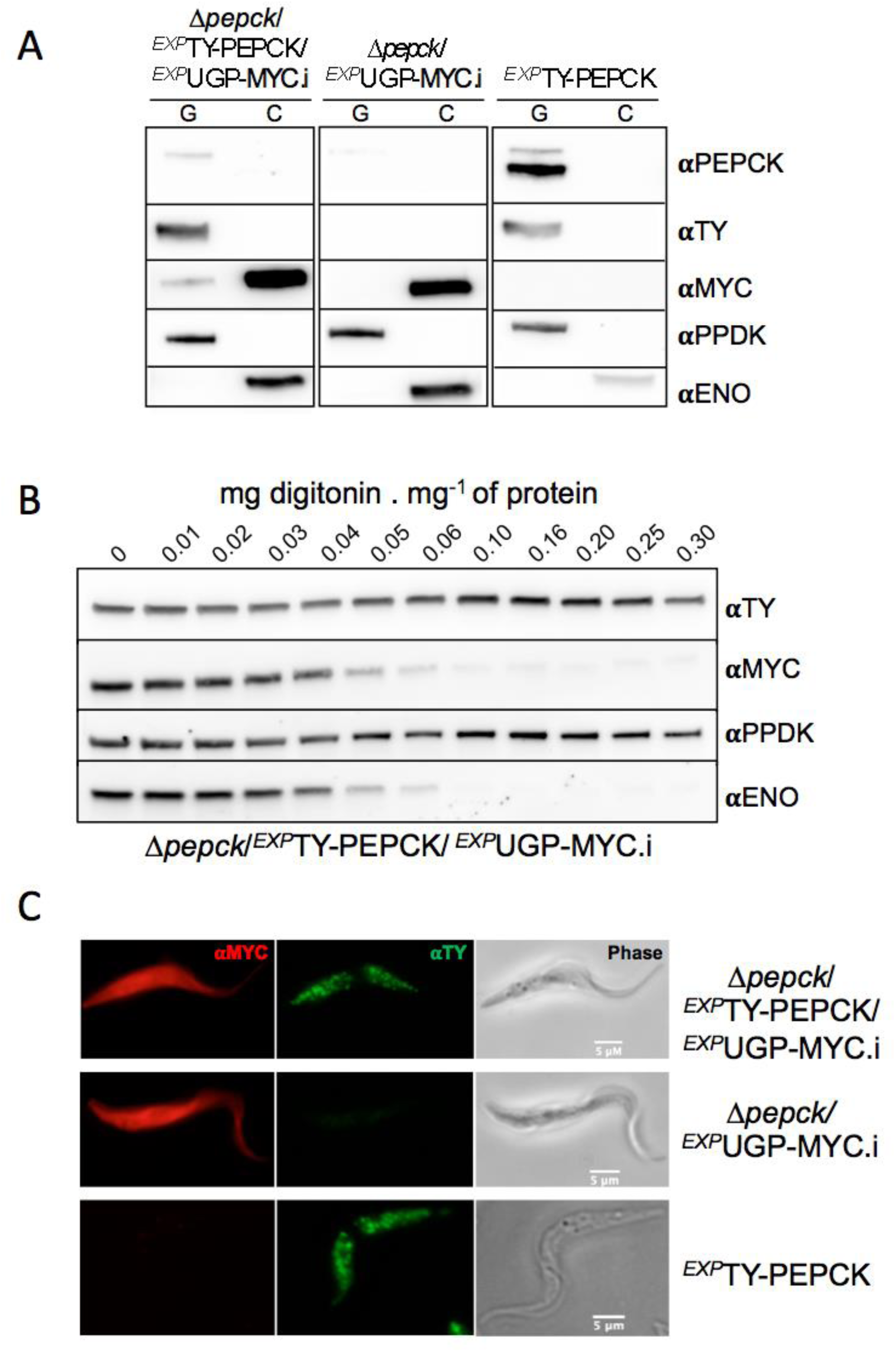
Analysis of UGP-MYC and TY-PEPCK tagged cell lines. Panel A shows western blot analysis of enriched glycosomal (G) and cytosolic (C) fractions obtained by differential centrifugation. The anti-PEPCK recognizes the tagged TY-PEPCK (αPEPCK) in addition to the native PEPCK. The samples were also analysed with the anti-TY (αTY) and anti-MYC (αMYC), as well as with glycosomal (αPPDK) and cytosolic (αENO) markers. Panel B shows digitonin titration analysis of the Δ*pepck*/*^EXP^*TY-PEPCK/*^EXP^*UGP-MYC.i cell line. Western blot analyses of pellets confirmed the glycosomal and cytosolic localisation of recombinant UGP-MYC with anti-MYC antibodies. Panel C shows the subcellular localisation of UGP-MYC and TY-PEPCK tagged proteins in the Δ*pepck*/*^EXP^*TY-PEPCK/*^EXP^*UGP-MYC, Δ*pepck*/*^EXP^*UGP-MYC and *^EXP^*TY-PEPCK cell lines by immunofluorescence microscopy analyses using rabbit anti-MYC (αMYC) and mouse anti-TY (αTY) antibodies.

**Fig. S2:**
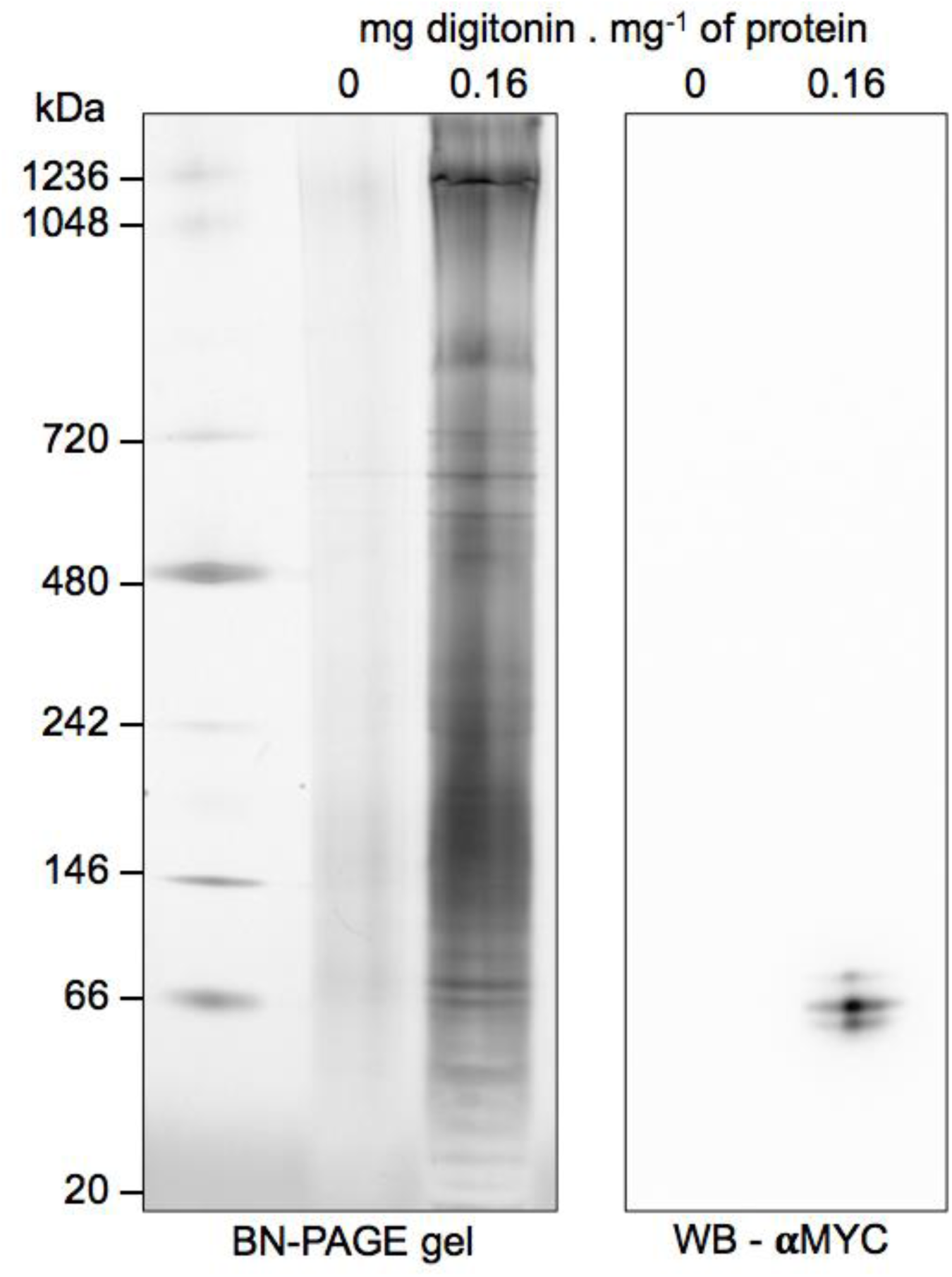
Analyse of UGP oligomerisation in native gel. The oligomeric state of tagged UGP-MYC was evaluated in the *^EXP^*UGP-MYC.i cell line. Supernatants obtained after digitonin treatment (0.16 mg of digitonin per mg of protein) or not (0) were analysed by BN-PAGE (left panel) and western blot (right panel) using the anti-MYC (αMYC) antibody.

**Fig. S3:**
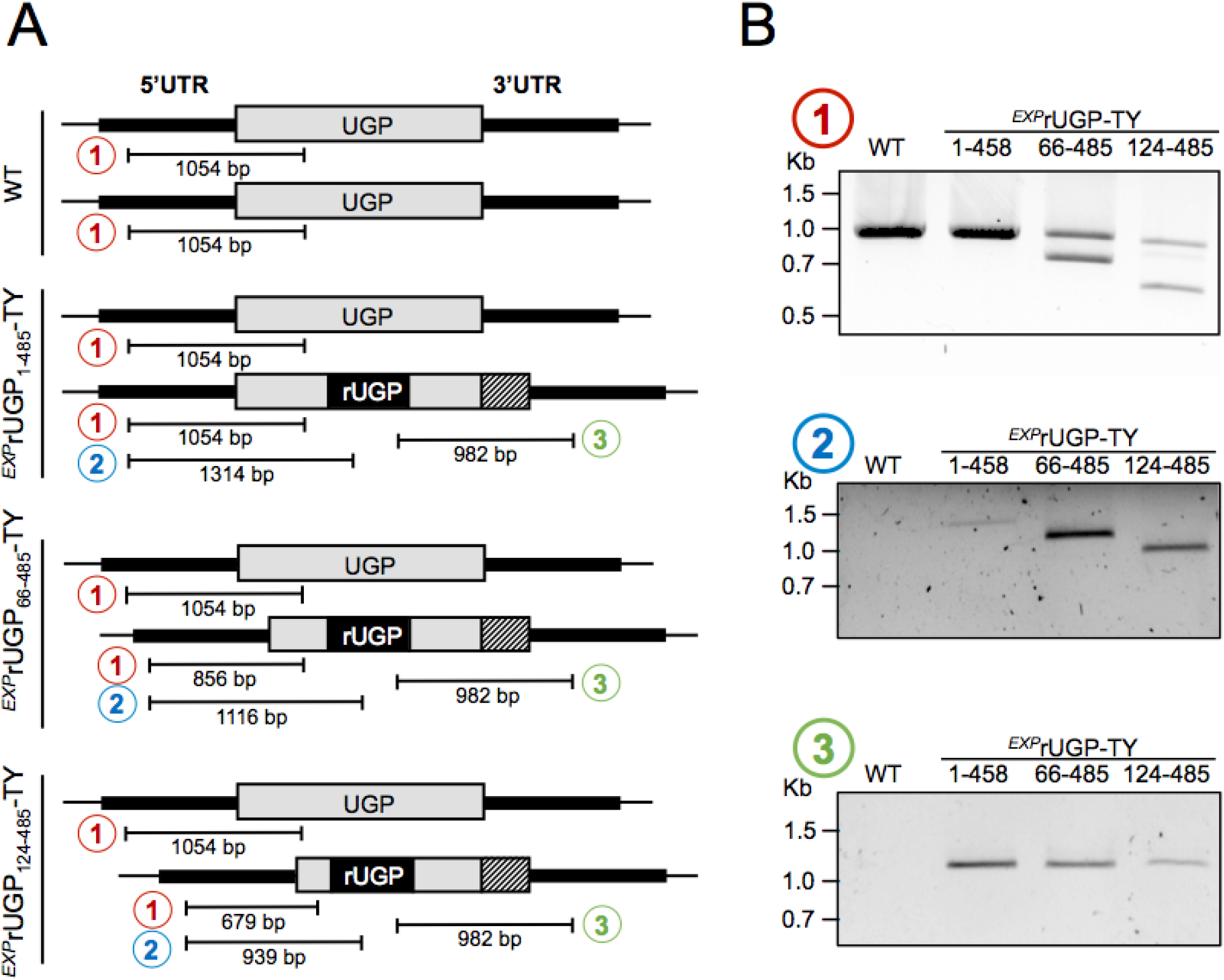
PCR analysis to confirm endo-tagging at the *UGP* locus. Panel A shows a schematic representation of the *UGP* native and tagged alleles in WT, *^EXP^*rUGP_1-485_-TY, *^EXP^*rUGP_66-485_-TY and *^EXP^*rUGP_164-485_-TY cell lines. The primers were designed to amplify products to differentiate between the native (1) and the recoded (2) tagged allele. The recoded region (rUGP) is highlighted by black boxes. PCR product number 3 confirms the TY-tagging at the C-terminal end. Panel B shows the PCR analysis of product 1, 2 and 3 using genomic DNA from parental cells as control.

**Fig. S4.**
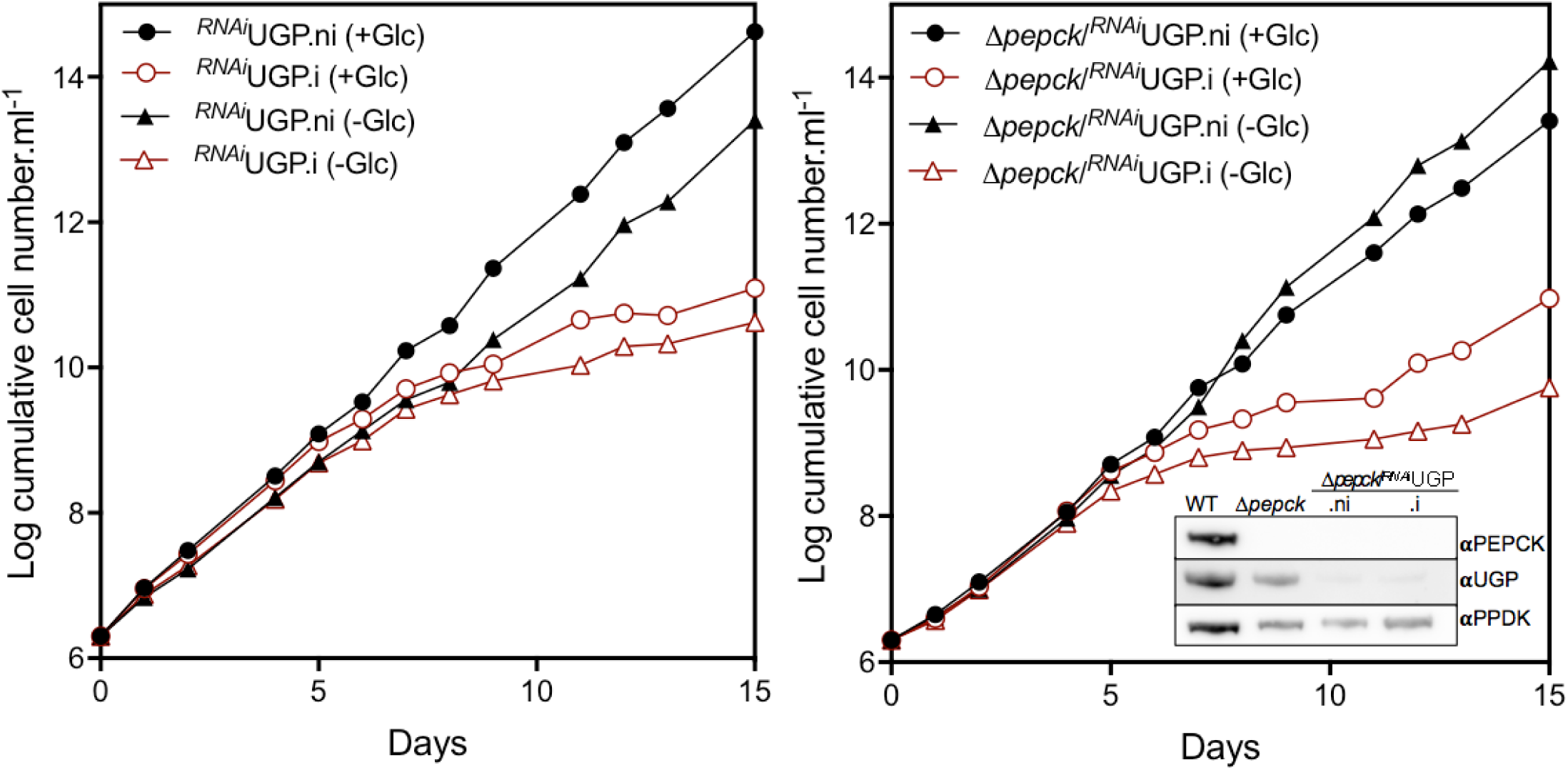
Functional analysis of *^RNAi^*UGP cell lines in presence or absence of glucose. Growth curves of the tetracycline-induced (.i) and non-induced (.ni) *^RNAi^*UGP and Δ*pepck*/*^RNAi^*UGP cell lines. The cells were grown in SDM79-GlcFree medium in the presence of 10 mM glucose (+Glc) or in the absence of glucose with 50 mM N-acetyl-glucosamine to inhibit the uptake of residual glucose (-Glu). The expression of PEPCK and UGP was monitored by western blot analysis (inset right panel) with the immune sera indicated in the right margin.

**Fig. S5.**
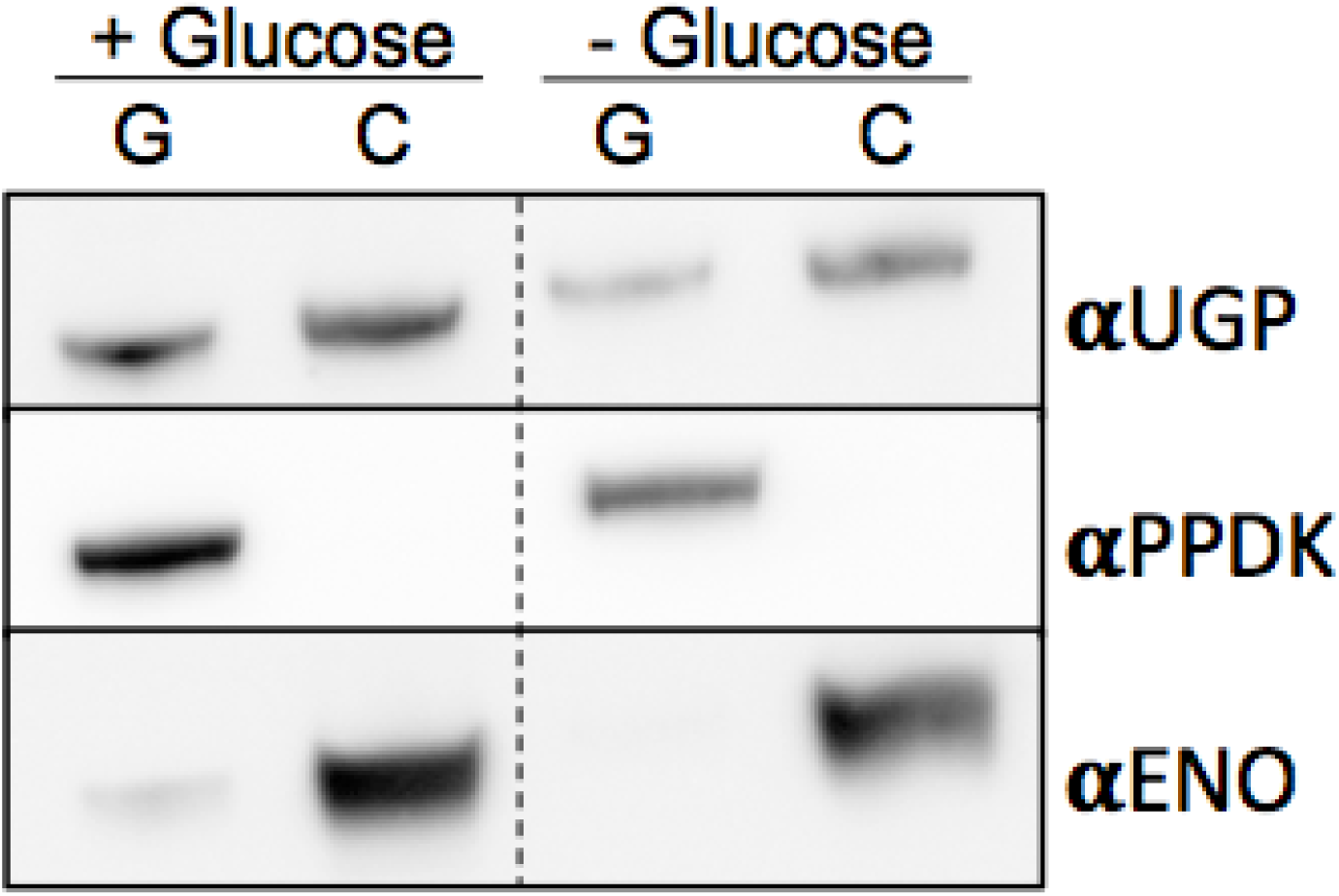
Subcellular localisation of UGP in the presence or the absence of glucose. The ratio glycosomal/cytosolic UGP was compared between the procyclic trypanosomes grown in the presence (+) or the absence (-) of glucose. The figure shows a western blot analysis of glycosomal (G) and cytosolic (C) fractions obtained after subcellular fractionation. Glycosomal (αPPDK) and cytosolic (αENO) markers are also shown.

**Fig. S6.**
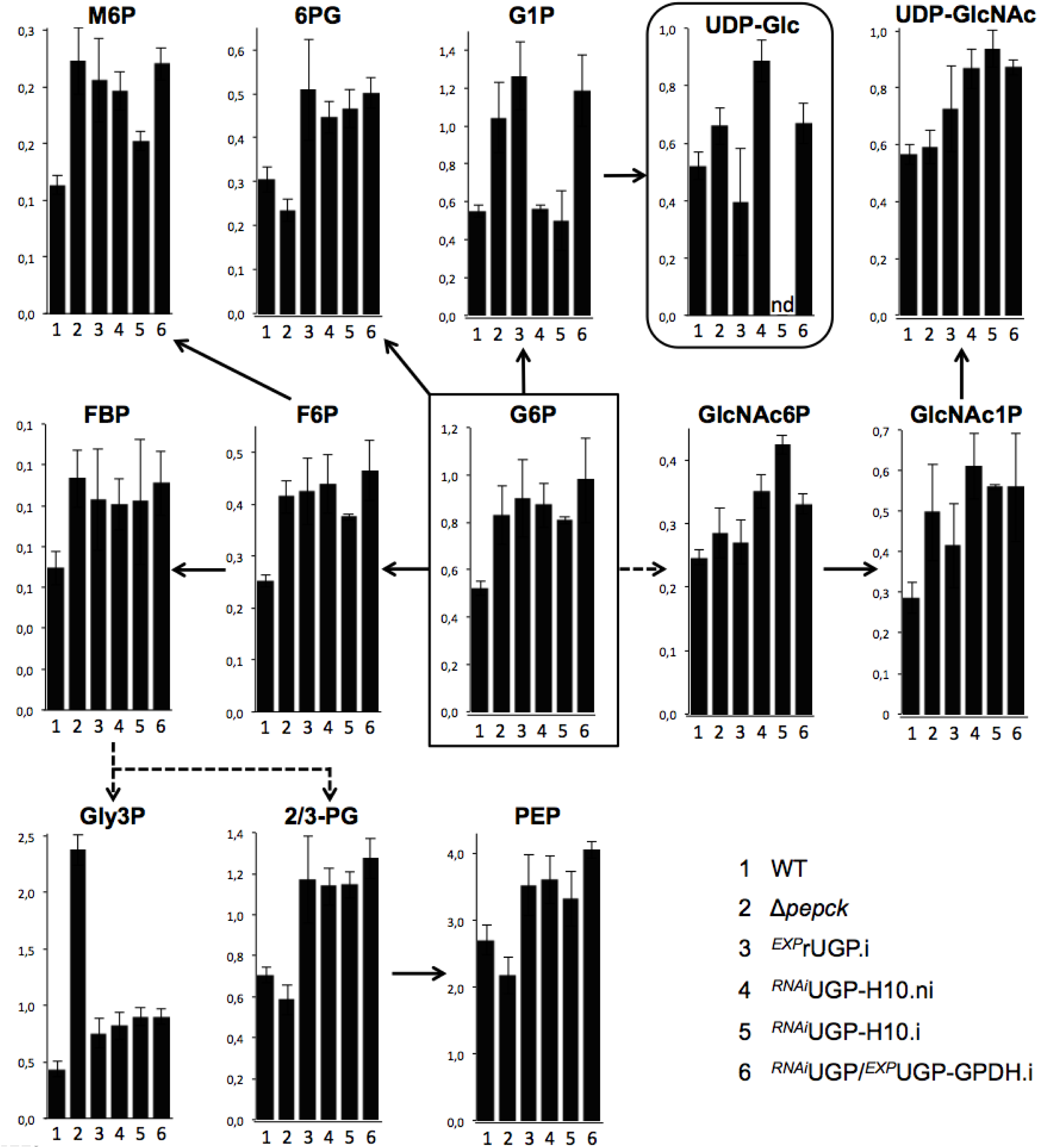
IC-HRMS analyses of intracellular metabolites collected from the indicated cell lines incubated in glucose-rich SDM79 medium.

**Fig. S7.**
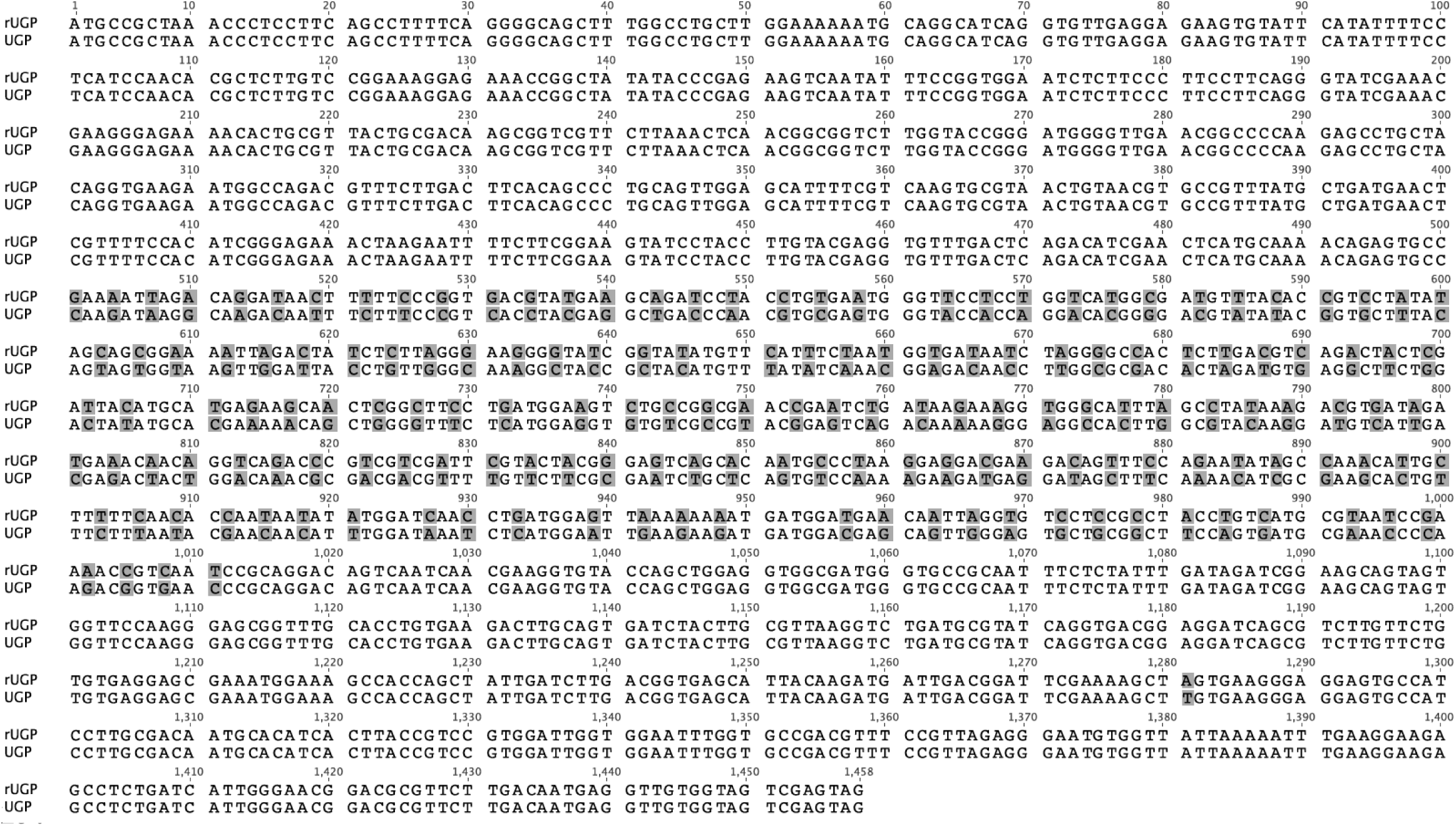
Sequence alignment of the recoded UGP sequence (rUGP) and native UGP. The residues modified to change the coding sequence without affecting the coded amino acid residue are shaded. For cloning purposes, the last nucleotide from restriction site HindIII at position 1276 was modified (shaded T in UGP and A in rUGP).

**Table S1.**
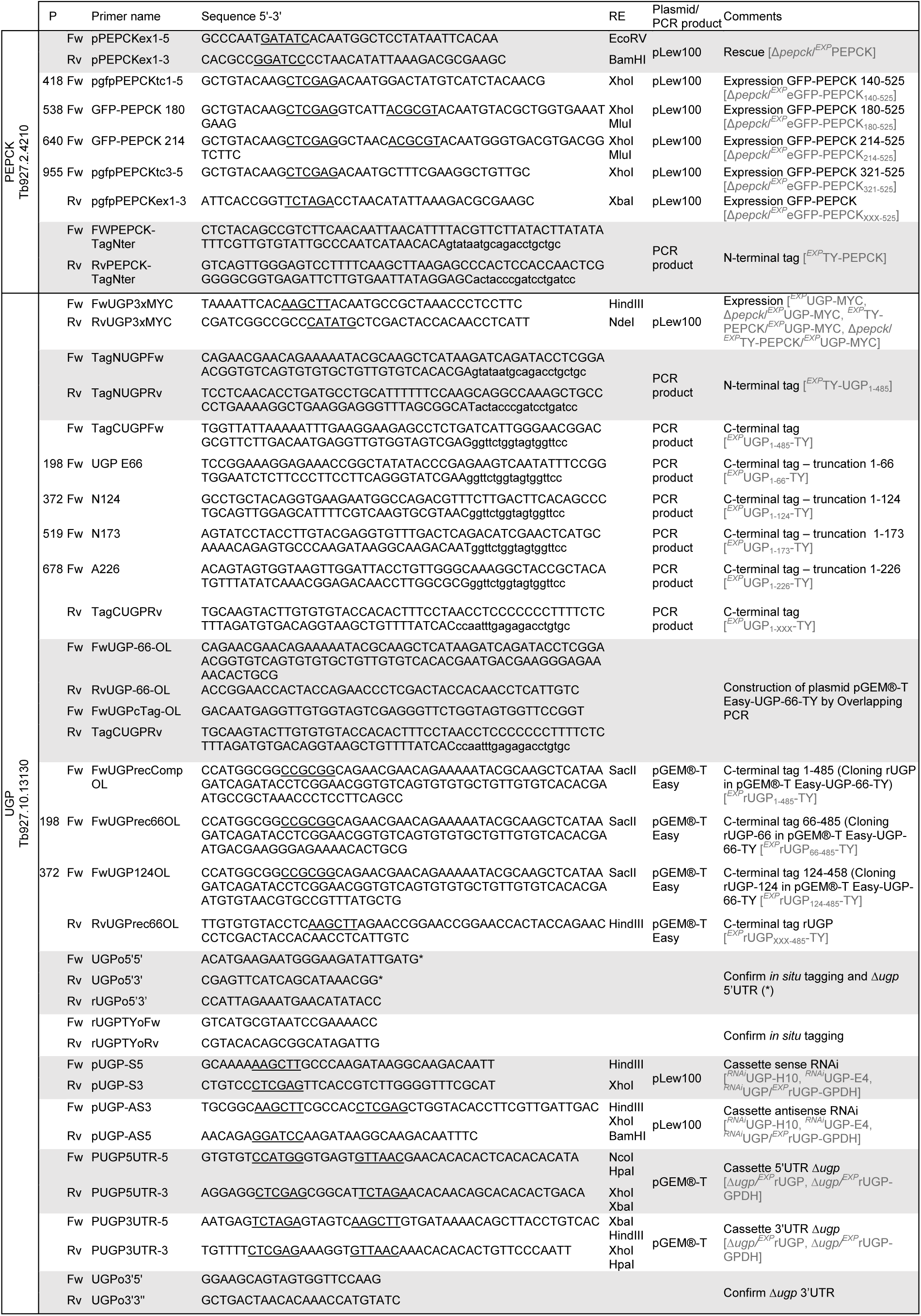
PCR primers sequences and features. P: Position in the gene; Fw: Forward primer; Rv: Reverse primer; RE: restriction sites (underlined). pPOT annealing sequences are shown in lower case; cell lines names are in box brackets.

**Table S2.**
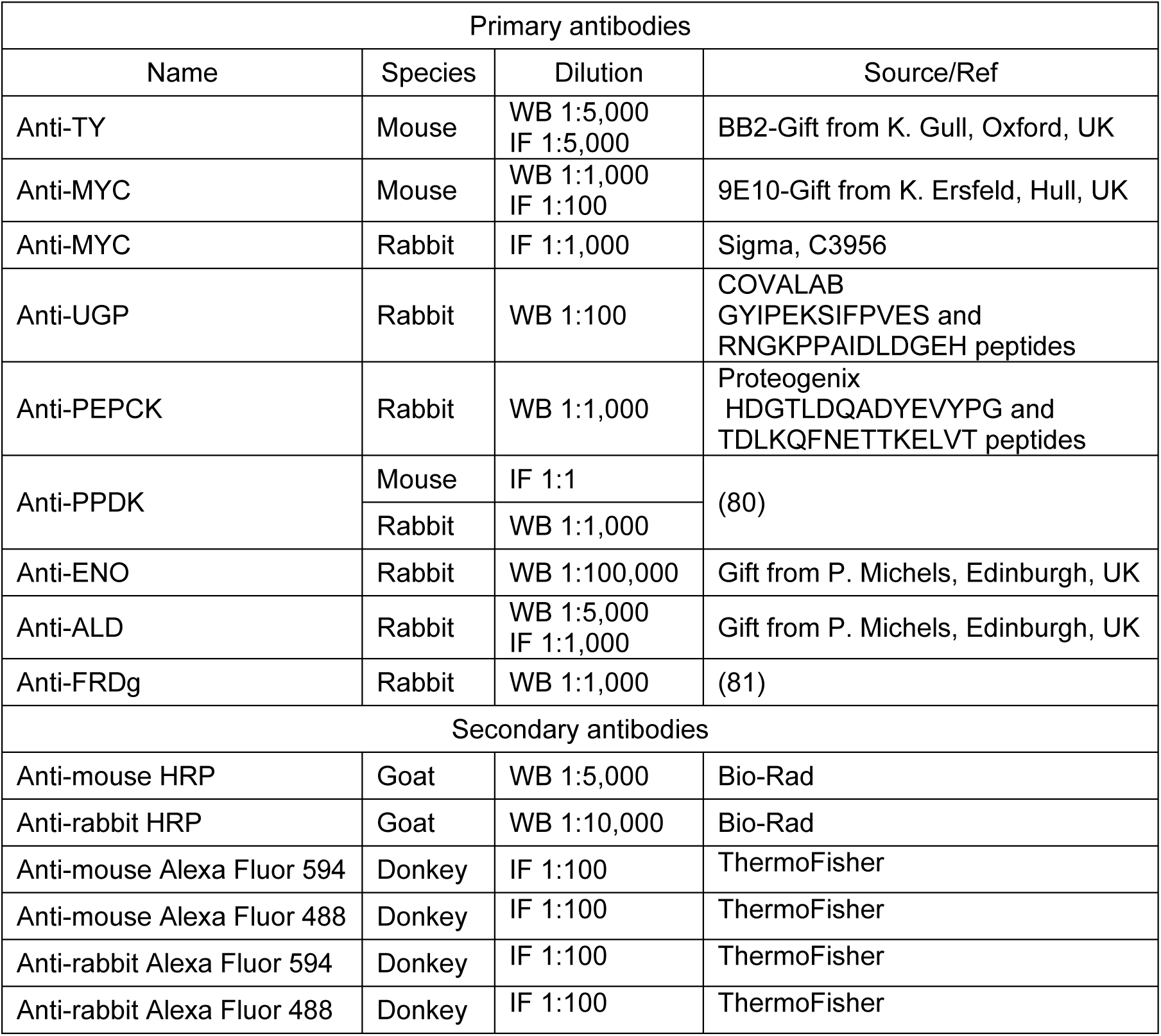
Antibodies used in this study.

## Acknowledgements

We thank Paul A. Michels (Edinburgh, Scotland) for providing us with the anti-enolase and anti-aldolase immune sera, as well as Keith Gull (Oxford, UK) and Klaus Ersfeld (Hull, UK) for providing us with the anti-TY (BB2) and anti-MYC (9E10) monoclonal antibodies, respectively. FB’s group was supported by the Université de Bordeaux, the Centre National de la Recherche Scientifique (CNRS), the ANR through the grants GLYCONOV (grant number ANR-15-CE15-0025-01) of the ANR-BLANC-2015 call and the French Government (Agence Nationale de la Recherche) Investissement d’Avenir programme, Laboratoire d’Excellence (LabEx) “French Parasitology Alliance For Health Care” (ANR-11-LABX-0024-PARAFRAP). MetaboHub-MetaToul (Metabolomics & Fluxomics facilities, Toulouse, France, http://www.metatoul.fr) is supported by the ANR grant MetaboHUB-ANR-11-INBS-0010.

